# Coupled beta and high-frequency oscillations emerge from synchronized bursting in a minimal model of the parkinsonian subthalamic nucleus

**DOI:** 10.64898/2026.03.30.715339

**Authors:** Hiba Sheheitli, Luke A. Johnson, Jing Wang, Joshua E. Aman, Jerrold L. Vitek

## Abstract

Local field potentials recorded from the subthalamic nucleus (STN) in Parkinson’s disease (PD) exhibit a distinctive multiscale spectral signature: exaggerated beta-band oscillations (13-30 Hz) coupled to high-frequency oscillations (HFOs, 200-400 Hz), with HFO amplitude being phase-locked to the beta cycle. This phase-amplitude coupling (PAC) has been identified as a promising biomarker of the parkinsonian state, yet no biophysical model has explained how it emerges, what determines the HFO frequency, or how HFOs can exist without beta modulation in the medicated STN. Here we show that a heterogeneous population of excitatory Izhikevich neurons with recurrent coupling produces three dynamical regimes: (i) asynchronous tonic firing, (ii) asynchronous bursting, in which neurons burst individually producing broadband HFO power but without coherent population-level PAC, and (iii) synchronous bursting, which gives rise to beta-HFO PAC. The regimes are governed by two biophysically interpretable parameters that capture complementary effects of dopamine depletion: one reflecting changes in intrinsic neuronal excitability, the other reflecting changes in synaptic coupling strength. The transition from asynchronous to synchronous bursting in this model captures the emergence of pathological STN neuronal activity in the parkinsonian state. HFO peak frequency varies continuously across the two-parameter landscape, suggesting a possible mechanism of the clinically observed shift from slow (200-300 Hz) to fast (300-400 Hz) HFOs between medication states. The character of the synchronization transition depends on baseline excitability, ranging from a sharp co-emergence of bursting and synchrony at low excitability to a decoupled transition at intermediate excitability, where the bursting fraction saturates while the population synchronization continues to increase with coupling. We also extend the model to include reciprocal coupling to an inhibitory population of globus pallidus externa (GPe)-like neurons and report a similar asynchronous-to-synchronous bursting transition and beta-HFO PAC emerging in the STN population, with the beta rhythm being set by the delay in the synaptic coupling between the two populations. The model generates testable predictions for future clinical and experimental studies, provides a numerical dissection of how mesoscopic LFP features map onto microscopic neuronal dynamics, and serves as a computational building block for future circuit-level models that may inform brain stimulation strategies tailored to the patient-specific dynamical state of the STN.

**Author summary:** In Parkinson’s disease, local field potentials (LFP) from the subthalamic nucleus (STN) contain two prominent rhythms: a slow beta rhythm (13-30 Hz) and fast oscillations (200-400 Hz). In the parkinsonian state, these rhythms become coupled, with fast oscillation amplitude varying systematically with beta phase, a relationship absent in the medicated state. We built a biophysical spiking neuron network model that captures two key effects of dopamine depletion on STN neuronal activity: changes in the intrinsic neuronal excitability and changes in synaptic coupling strength. The model produces fast oscillations from rapid intraburst firing, while the slow beta rhythm and its coupling to fast oscillations emerge with the onset of synchronized bursting across the population. Importantly, the frequency of the fast oscillations shifts continuously depending on both parameters, explaining a puzzling clinical observation that these oscillations change frequency between medication states. The model also reproduces the modulation pattern in the spike-triggered average of HFO envelope amplitude reported in patient recordings, confirming consistency with single-unit observations as well as LFP-level spectral features. By mapping how multi-timescale LFP spectral features relate to the dynamical regime of the underlying neuronal population, this work offers a framework for brain stimulation strategies informed by patient-specific dynamical states.

## Introduction

Exaggerated beta-band (13-30 Hz) oscillatory activity in the subthalamic nucleus (STN) is a hallmark electrophysiological signature of Parkinson’s disease (PD) and has been reported to correlate with motor symptom severity [1–3]. Alongside beta oscillations, high-frequency oscillations (HFOs) above 200 Hz have been reported in STN local field potentials (LFPs) in both the parkinsonian (OFF-medication) state and the medicated (ON-medication) state [4,5]. The clinical literature has categorized STN HFOs into a slow component (sHFO, 200-300 Hz) and a fast component (fHFO, 300-400 Hz), with dopaminergic medication causing a characteristic power shift from sHFOs toward fHFOs [5,6]. The ratio of sHFO to fHFO power correlates with clinical motor scores and has been proposed as a neurophysiological marker of the dopaminergic and motor state [5]. Notably, this HFO power ratio correlated with clinical scores but not with beta peak power, yet a combined parameter incorporating both provided the strongest clinical correlation [5], suggesting that more than one physiological factor may underlie the pathological state.

Critically, what distinguishes the pathological state is not the mere presence of HFO power but the strong modulation of HFO amplitude by the phase of beta oscillations, with the strength of this phase-amplitude coupling (PAC) in the STN correlating with bradykinesia and rigidity severity [7,8]. This is in contrast to what is observed in the non-parkinsonian medicated STN, where HFOs may be present but are movement-modulated, and as such described as functionally prokinetic or “liberated” from beta modulation [7]. Meidahl et al. [9] reported that this PAC reflects the synchronization of single-unit spiking to network beta oscillations: in the presence of PAC, the spike-triggered average (STA) of the LFP shows clear HFO-frequency oscillatory structure and inter-spike interval (ISI) distributions peak at HFO frequencies. These findings suggest that PAC captures a fundamental aspect of pathological neuronal spiking patterns rather than a spectral epiphenomenon.

This body of clinical evidence motivates the effort to develop mechanistic models that can concurrently capture the beta and HFO rhythms and provide explanatory insight into the relationship between LFP spectral features and the underlying neuronal spiking dynamics. Three specific questions remain open. First, what biophysical mechanism could generate beta-modulated HFOs in a population of STN-like neurons? Second, what determines the HFO frequency, and why does it shift between medication states? Third, how can HFOs exist without beta modulation in the medicated state, and what kind of transition in neuronal spiking dynamics could underlie the emergence of the pathological beta-HFO PAC?

Previous computational models of the basal ganglia have focused primarily on beta oscillation generation through multi-region circuit models involving basal ganglia-thalamocortical loops [10–13]. These models successfully reproduce beta-band synchronization but do not generate HFOs or address cross-frequency coupling, and in general, much less attention has been given to HFO dynamics and its possible relationship to slower rhythms. A recent exception is Yu et al. [14], who developed a primary motor cortex-basal ganglia-thalamus model that reproduces beta-broad gamma PAC in the 100-200 Hz range through multi-region interactions, but does not address the 200-400 Hz HFOs recorded in the STN or the mechanism that determines HFO frequency. From an alternative modeling perspective, exact mean-field models of quadratic integrate-and-fire (QIF) neurons with a slow adaptive current produce beta-nested fast oscillations [15–17], also known as mixed-mode oscillations [18], indicating that such multiscale rhythms arise generically from a separation of timescales. However, the reset potential is absent from the reduced mean-field equations, so the formulation cannot distinguish a population of intrinsic bursters from one of tonic-firing neurons, the distinction that is central to the parkinsonian STN, where the pathological state is marked by increased bursting and burst synchrony across the population [3]. This motivates us to examine the dynamics at the spiking neuron population level, such that we can capture the neuron-level propensity to bursting and map the emergent population-level multiscale oscillations onto both the underlying spiking dynamics and the corresponding LFP spectral features. Sanders [19] demonstrated that a population of stochastic burst generators with prescribed firing rate statistics produces PAC resembling that observed in parkinsonian animals, providing an important proof of concept that synchronized bursting can give rise to PAC. However, their model prescribes burst parameters as inputs rather than deriving them from biophysical dynamics, and does not address what controls HFO frequency or the transition from the healthy to the pathological state.

Here we construct a minimal single-population model using excitatory Izhikevich (adaptive QIF) spiking neurons [20] with slow adaptation and recurrent coupling, a framework that models both intrinsic and synaptic changes associated with dopamine depletion in a computationally efficient setting. We map the full two-parameter landscape defined by the median reset potential (*c̅*, controlling intrinsic excitability) and synaptic coupling strength (J). These parameters have clear dopaminergic substrates: *c̅* reflects dopamine-dependent modulation of intrinsic membrane currents that govern the tonic-to-burster transition in STN neurons [21,22], while J reflects changes in effective excitatory synaptic drive within the STN. The use of a single excitatory population with recurrent coupling is motivated by experimental evidence that STN neurons produce spontaneous synchronized burst firing through glutamatergic axon collaterals, even when isolated from all external inputs [23], although functional recurrent excitation between STN neurons is reported to be weak or undetectable in other studies [24–26]. Hence, we interpret J as an effective coupling strength rather than a literal count of recurrent STN synapses. The stronger coupling associated with the parkinsonian state can then reflect increased synchronizing coupling at the network level, for instance through an STN-GPe loop or shared synaptic input, rather than increased recurrent excitation within the STN.

We show that this model produces three dynamical regimes spanning tonic firing, asynchronous bursting, and synchronous bursting. The clinically relevant transition is from asynchronous bursting (HFO power without PAC) to synchronous bursting (beta-HFO PAC), corresponding to the medicated/ON-to-parkinsonian/OFF state change. The path to synchronization depends on baseline excitability: when neurons fire tonically at baseline, increasing coupling drives them directly into synchronized bursting, whereas when neurons are intrinsic bursters, HFO power is already present without PAC at low coupling, and pathological (i.e., abnormally elevated) beta-HFO coupling emerges when their bursting becomes coordinated and time-locked across the population. The HFO frequency emerges from the aggregate of intraburst firing rates across the population and varies continuously across the (c̅, J) landscape, providing a mechanistic explanation for the clinically observed sHFO/fHFO shift between medication states. The model also reproduces the modulation of the STA of HFO envelope amplitude reported by Meidahl et al. [9], confirming consistency between the synchronization mechanism and single-unit recordings. This work provides a numerical dissection of how multi-timescale mesoscopic LFP spectral features map onto the microscopic neuronal operating regime, and serves as a building block for future circuit models incorporating the larger basal ganglia-thalamocortical network. The framework may inform DBS strategies that target personalized dynamical regimes (i.e., modes of collective neuronal activity) rather than relying on generic one-dimensional spectral biomarker thresholds such as beta amplitude.

## Results

### Single-neuron phenomenology: the tonic-to-burster transition

The single-neuron dynamics of the Izhikevich model (see Methods) provides the mechanistic substrate for the population-level phenomena investigated in this study. The reset potential c controls the depth of the after-spike hyperpolarization and thereby determines whether a neuron fires tonically or in bursts (Fig 1A). At low values (c = -65 mV), the deep reset suppresses the slow recovery variable sufficiently to produce regular single spikes. As c increases past approximately -57 mV, the less negative reset places the membrane potential closer to threshold after each spike, where the quadratic depolarizing drive is sufficient to reach threshold again rapidly, producing a burst of spikes. Each spike increments the adaptation variable u by a fixed amount d, and the burst terminates when u accumulates enough to prevent the membrane from reaching threshold. The result is a separation of timescales intrinsic to each neuron: fast intraburst spiking at rates in the HFO range (200-400 Hz) and slow interburst intervals at rates in the beta range (13-30 Hz), as revealed by the bimodal ISI distributions in Fig 1B.

**Fig 1.**
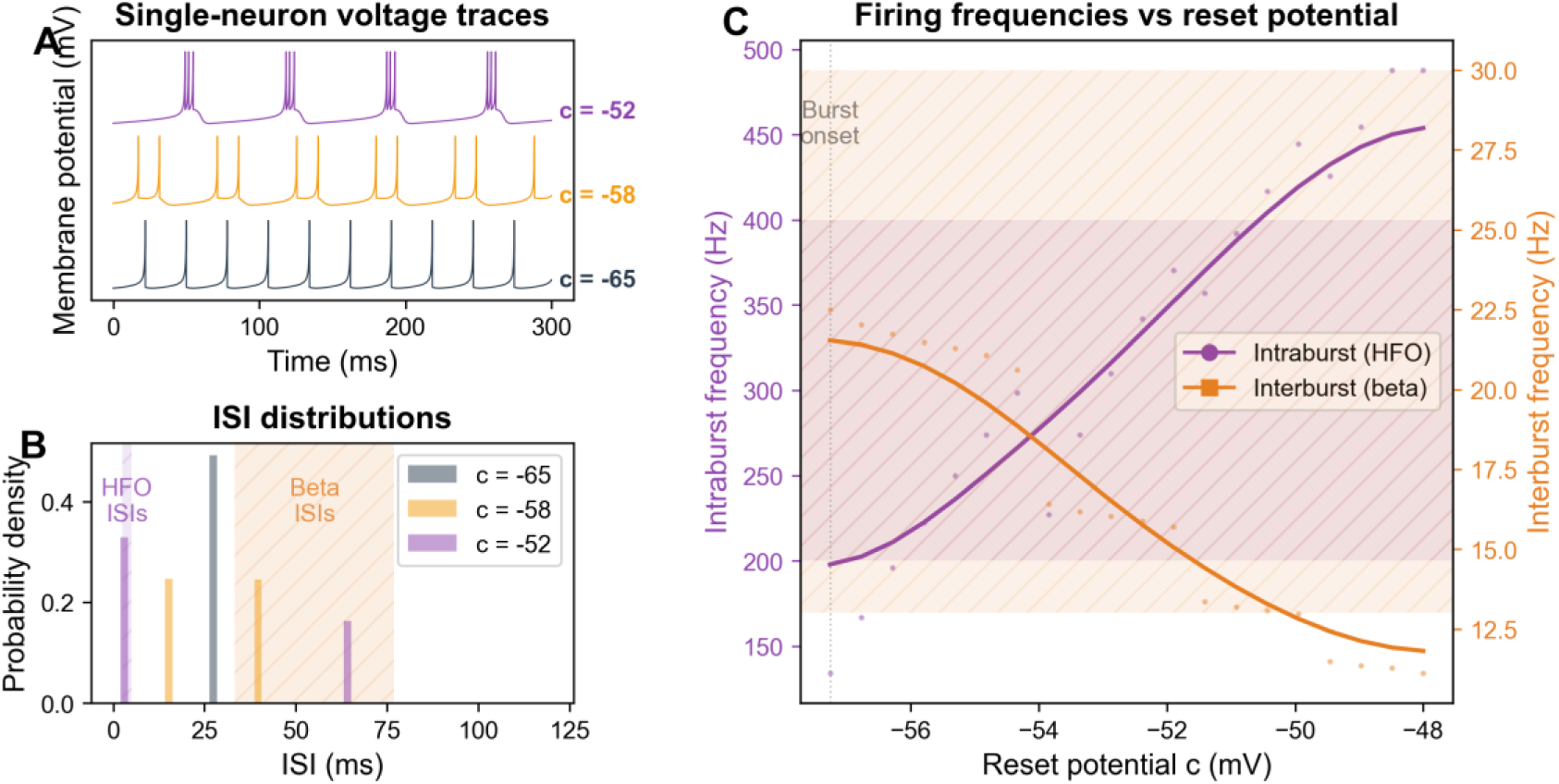
Single-neuron phenomenology. (A) Voltage traces at representative c values: tonic (c = -65 mV), bursting with short bursts (c = -58 mV), robust bursting (c = -52 mV). (B) ISI probability density distributions at representative c values; shaded bands mark the canonical beta (13-30 Hz, orange) and HFO (200-400 Hz, purple) frequency ranges for reference. (C) Intraburst (HFO, purple) and interburst (beta, orange) frequencies in the model as a function of reset potential c. Dots show values at each c; solid lines are smoothed trends.

Critically, both timescales depend continuously on c (Fig 1C). The neuron’s adaptation variable sets the interburst (beta) timescale phenomenologically and is not intended to represent any specific spike frequency adaptation reported in STN neurons which might operate on different timescales. This inverse relationship between the two characteristic frequencies of the bursting neuron, encoded in a single biophysical parameter, establishes the foundation for the continuous HFO frequency landscape at the population level and provides the mechanism by which changes in intrinsic excitability can shift HFO frequency between medication states.

### Network dynamics: from tonic firing to asynchronous and synchronous bursting

The character of the synchronization transition depends fundamentally on how the population is distributed relative to the single-neuron tonic-to-bursting boundary. We present three representative values of median reset potential *c̅* that illustrate qualitatively distinct dynamical scenarios.

At low median intrinsic excitability (*c̅* = -64 mV, Fig 2), the bulk of the population lies below the bursting threshold: at weak coupling, neurons fire asynchronous tonic spikes and the LFP fluctuates irregularly, with a broad spectral peak near the mean tonic firing rate (∼35 Hz) but no sharp oscillatory structure and no HFO power. Recurrent excitatory coupling acts as an effective depolarizing drive that shifts neurons toward the tonic-to-bursting boundary. Since barely any neurons burst intrinsically at this *c̅* value, the onset of bursting requires a critical level of recurrent input, and once that threshold is crossed, the same coupling that induces bursting also aligns burst timing across the population. The result is a single, relatively sharp transition in which bursting and synchronization co-emerge: HFO power, beta oscillations, and beta-HFO PAC all appear together at the synchronization boundary, with no intermediate regime of asynchronous bursting.

**Fig 2.**
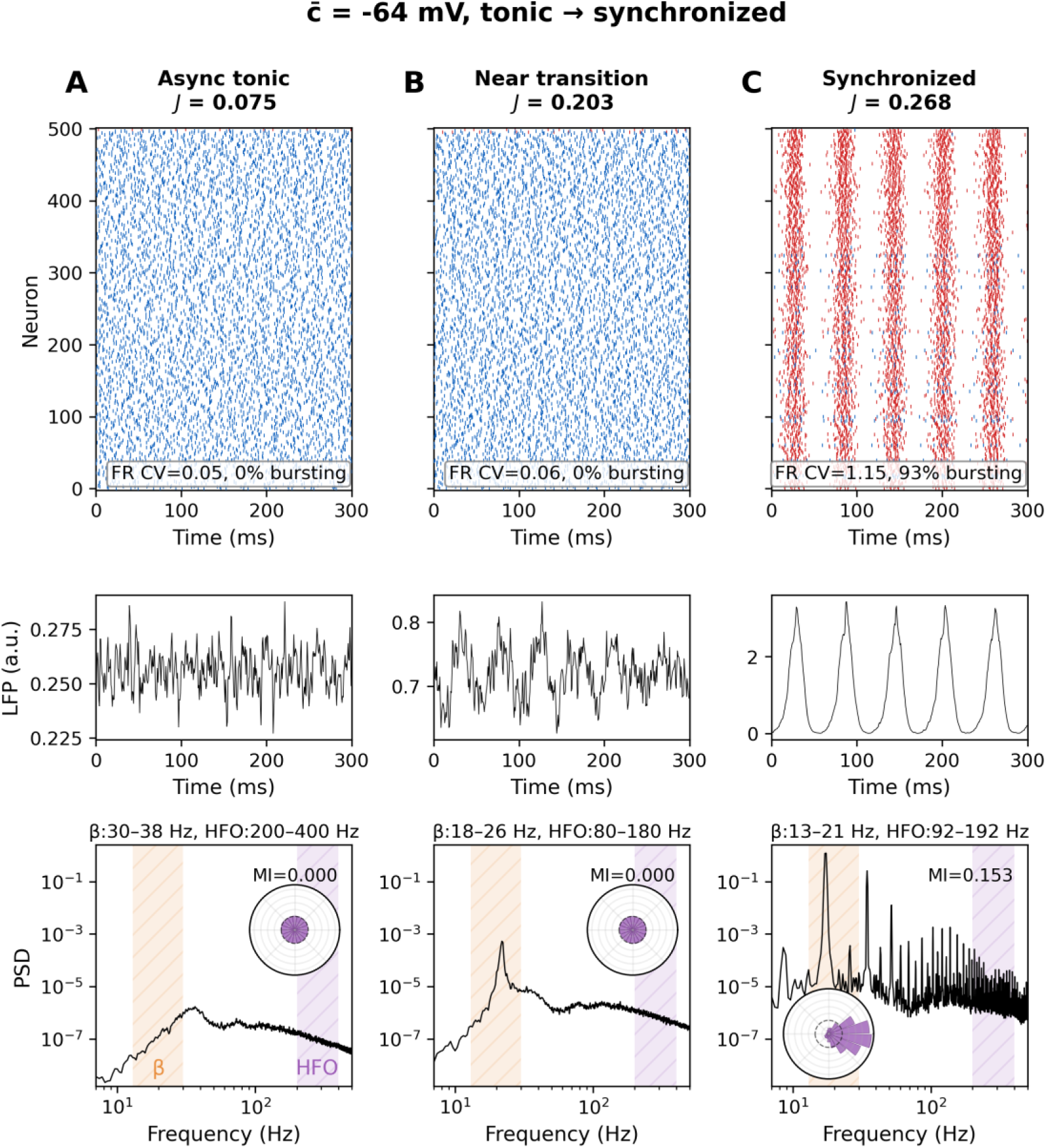
Synchronization at low median intrinsic excitability (*c̅* = -64 mV). Three panels showing progression from asynchronous tonic firing (A, J = 0.075) through the synchronization transition (B, J = 0.203) to synchronous bursting (C, J = 0.268). Each panel contains a raster plot (top row), synthetic LFP trace (middle row), and power spectral density of the LFP on log-log scale (bottom row). Rasters show 500 neurons evenly sampled from the full population of 5000, ordered by reset potential. In the raster plots, blue dots denote spikes from neurons classified as tonic and red dots denote spikes from neurons classified as bursting (see Methods); the text box reports the coefficient of variation (CV) of the mean firing rate (FR), used as a proxy for synchrony; see Methods) and the percentage of bursting neurons. In the PSD panels, shaded bands mark the canonical beta (13-30 Hz, orange) and HFO (200-400 Hz, purple) frequency ranges for visual reference. The text above each PSD panel reports the adaptive frequency bands used for PAC computation at that parameter point (detected slow peak ± 4 Hz for the phase band, detected HFO peak ± 50 Hz for the amplitude band; see Methods). The polar inset shows the mean HFO amplitude as a function of beta phase, with the corresponding Tort modulation index (MI) that quantifies the strength of PAC. For the non-zero MI value, the surrogate test gives z = 4.7, p < 0.005 (panel C).

At intermediate median intrinsic excitability (*c̅* = -58 mV, Fig 3), the Lorentzian spread of reset potentials places the population astride the single-neuron tonic to bursting boundary: approximately 37% of neurons are intrinsic bursters at baseline, while the remainder fire tonically. As coupling strength increases, beta-band power in the LFP increases while the bursting fraction and PAC remain near baseline (Fig 3A-B). At higher coupling, more neurons are recruited into bursting (Fig 3C), and the population locks together, producing strong beta oscillations and beta-HFO PAC (Fig 3D). The bursting fraction saturates while the population synchronization continues to increase with coupling, so recruitment and synchronization are separated in coupling strength, as illustrated in Fig 5.

**Fig 3.**
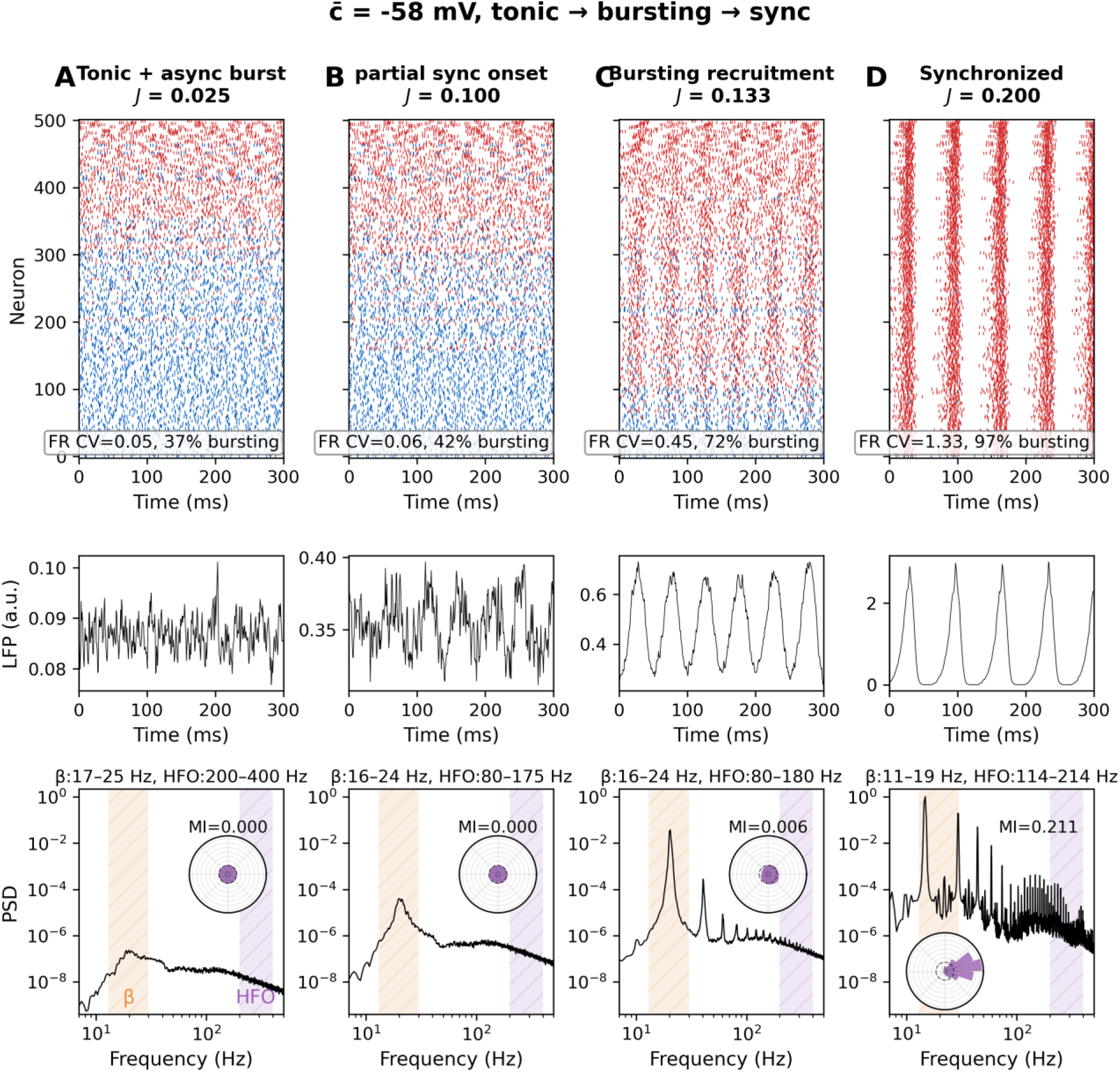
Synchronization at intermediate excitability (*c̅* = -58 mV). Four panels showing the transition from tonic-dominated asynchronous activity (A, J = 0.025) through increased beta-band power (B, J = 0.100) and recruitment of tonic neurons into bursting with partial synchrony (C, J = 0.133), to synchronous bursting (D, J = 0.200). Format as in Fig 2. For the non-zero MI values, the surrogate test gives z = 15.9, p < 0.005 (panel C) and z = 4.9, p < 0.005 (panel D).

At high median intrinsic excitability (*c̅* = -53 mV, Fig 4), nearly all neurons are intrinsic bursters even without any coupling. Each neuron independently produces HFO-frequency intraburst firing, but since burst onsets are uncorrelated in time across the population, the aggregate LFP shows an HFO power peak but without a sharp beta peak or beta-HFO PAC. This asynchronous bursting regime is the model’s counterpart of the medicated STN, where HFOs have been reported without pathological beta modulation [7,27]. As coupling increases, shared synaptic input progressively correlates burst timing across neurons. The transition to synchrony is purely a collective phenomenon: no change in individual neuronal firing mode is required, only the temporal alignment of pre-existing bursts. Once synchronized, the population LFP acquires a sharp beta peak at the common interburst frequency and a strong PAC emerges, that is, HFO amplitude becomes phase-locked to the beta cycle.

**Fig 4.**
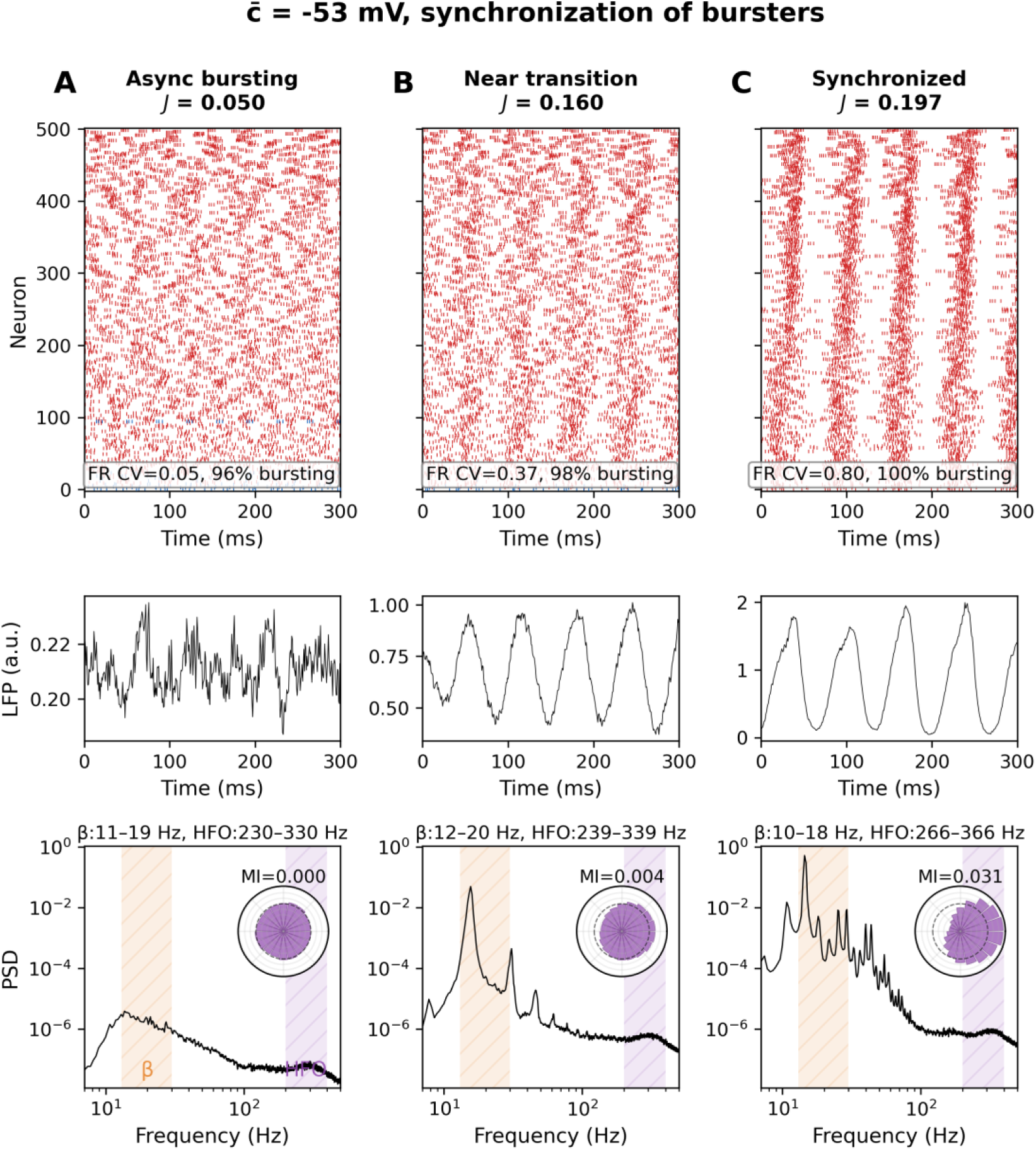
Synchronization at high excitability (*c̅* = -53 mV). Three panels showing progression from asynchronous bursting (A, J = 0.050) through partial synchrony near-transition (B, J = 0.160) to synchronous bursting (C, J = 0.197). Format as in Fig 2. For the non-zero MI values, the surrogate test gives z = 18.3, p < 0.005 (panel B) and z = 11.6, p < 0.005 (panel C).

These three scenarios form a continuum governed by the fraction of neurons that are intrinsic bursters at baseline. When this fraction is near zero (low *c̅*), coupling must both induce bursting and synchronize it, and the two processes co-occur because the recurrent excitation that pushes neurons past the bursting threshold simultaneously correlates their activity. When the fraction is intermediate, the bursting fraction saturates while the population synchronization continues to increase with coupling, so recruitment and synchronization are separated in coupling strength. When the fraction is near unity (high *c̅*), recruitment is unnecessary and the transition involves only synchronization. Four additional supporting figures, S1 through S4, trace this continuum at additional intermediate *c̅* values: at *c̅* = -62 mV (Fig S1) and *c̅* = -60 mV (Fig S2), where baseline bursting fractions are low and the transition resembles the co-emergent pattern of Fig 2; and at *c̅* = -56 mV (Fig S3) and *c̅* = -54 mV (Fig S4), where baseline bursting fractions are high and the transition is dominated by synchronization of pre-existing bursters as in Fig 4.

Fig 5 provides quantitative confirmation of this transition typology by plotting the co-evolution of the mean firing rate CV (coefficient of variation of the population firing rate; see Methods) and the fraction of bursting neurons as a function of coupling strength at the three representative c̅ values. At low excitability (Fig 5A), essentially no neurons burst at baseline (0%); bursting appears only as recurrent excitation recruits non-bursting neurons, so the bursting fraction and the mean firing rate CV rise together in a single sharp transition, consistent with the adaptation-driven population bursting analyzed for spiking networks in [17]. At intermediate excitability (Fig 5B), a substantial fraction already bursts at baseline (37%) and the bursting fraction saturates at weaker coupling than the full synchronization transition, so the two curves separate. At high excitability (Fig 5C), most neurons are already intrinsic bursters at baseline (96%), each acting as an oscillator at its interburst frequency; the bursting fraction is saturated and the transition reflects only whether coupling synchronizes them.

**Fig 5.**
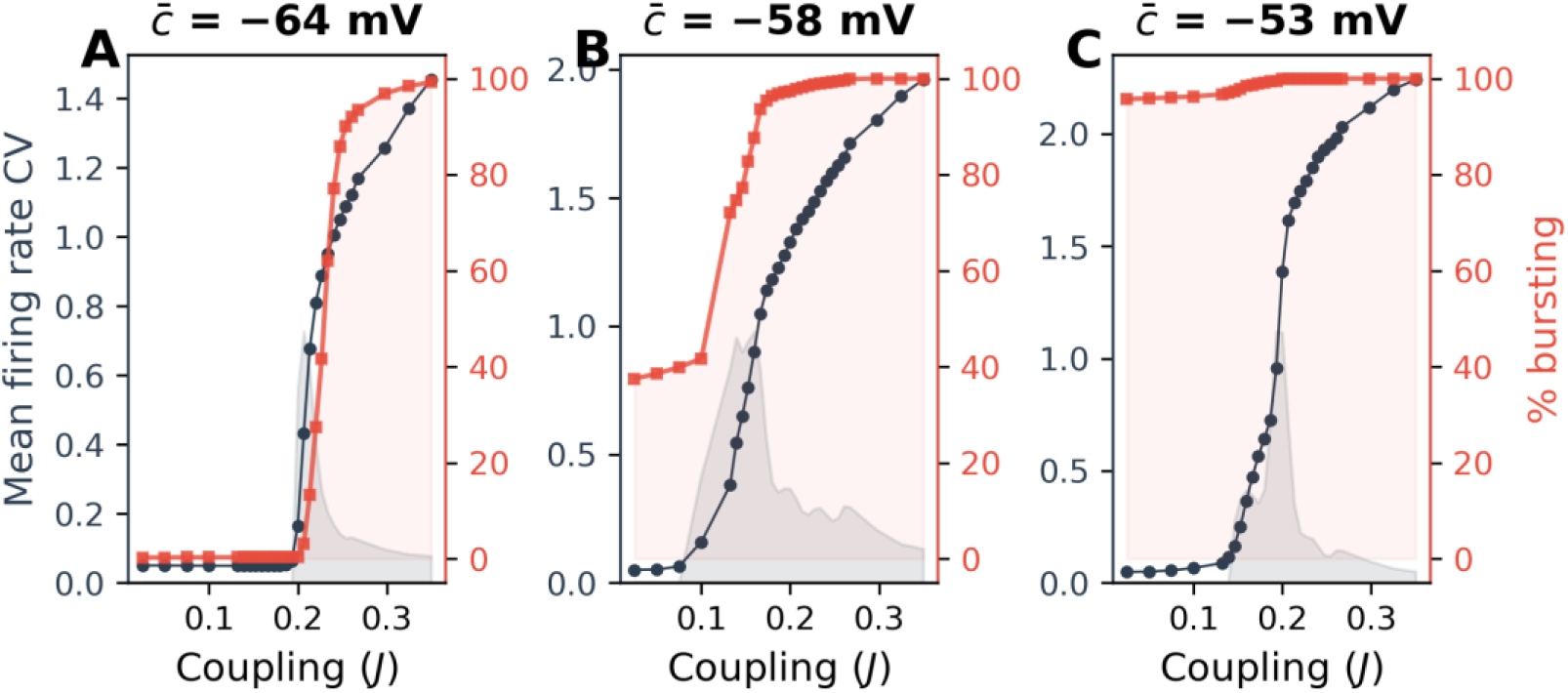
Decoupling of bursting and synchronization transitions. Each panel shows mean firing rate CV (dark line, left axis; a proxy for synchrony), percent bursting neurons (red line with squares, right axis), and the derivative of the CV curve (gray shading, arbitrary units) highlighting the steepness of the synchronization transition, as a function of coupling strength J. (A) Co-occurring transitions at c̅ = -64 mV: bursting fraction 0% at baseline. (B) Decoupled transition at c̅ = -58 mV: bursting fraction 37% at baseline; the fraction saturates while the CV continues to increase with coupling. (C) Synchronization-only transition at c̅ = -53 mV: bursting fraction already 96% at baseline.

### The excitability-coupling phase diagram

The full (c̅, J) parameter space is mapped in Fig 6, where synchrony (measured by the mean firing rate CV), relative beta (13-30 Hz) power, PAC modulation index, and HFO center frequency are displayed as heatmaps across the 2D parameter grid.

**Fig 6.**
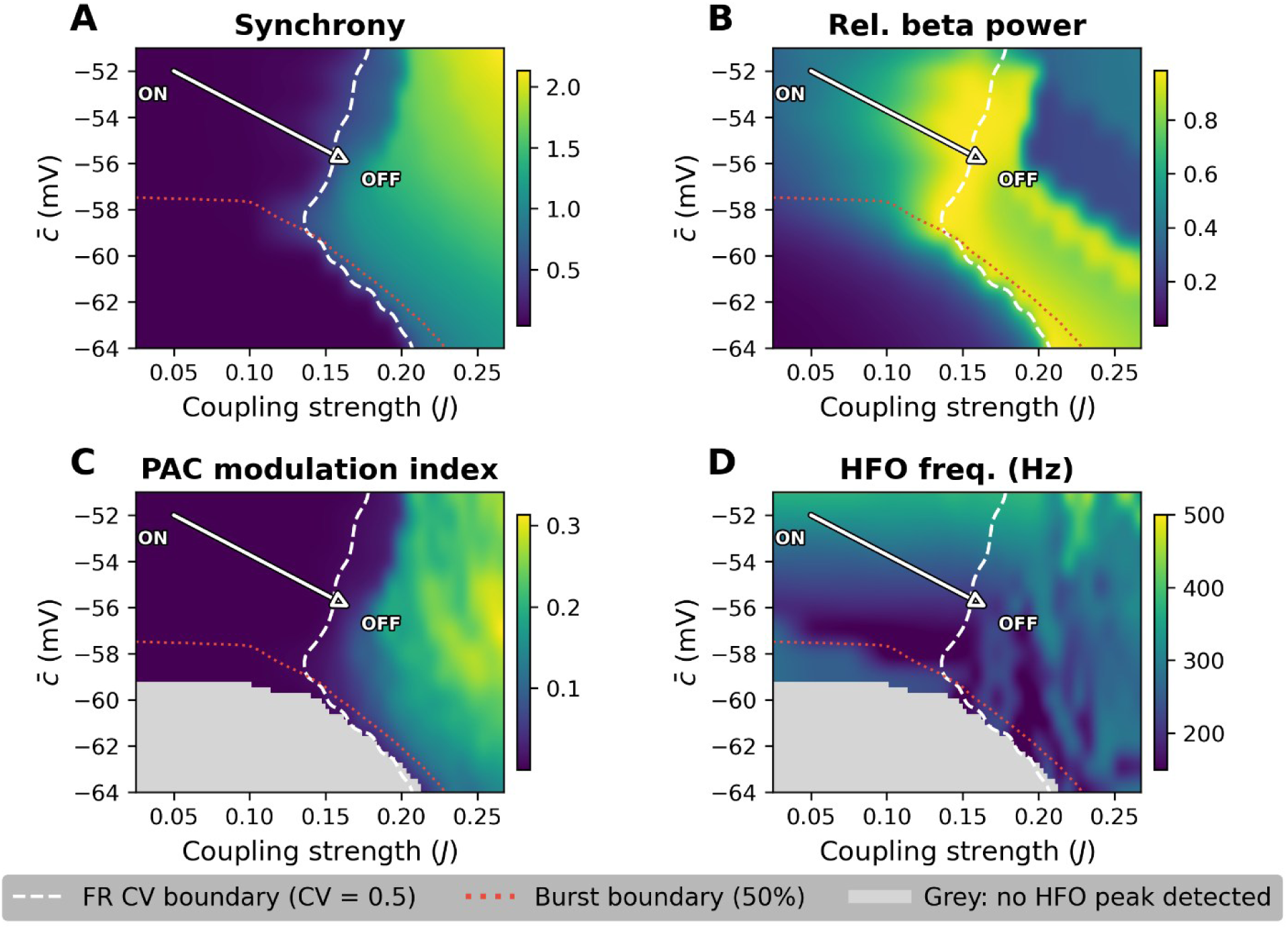
(*c̅*, J) phase diagram. (A) Synchrony, the mean firing rate CV (see Methods). (B) Relative beta power, the fraction of total spectral power in the 13-30 Hz band. (C) PAC modulation index (see Methods). (D) HFO center frequency (see Methods). Dashed white line: synchronization boundary (CV = 0.5). Dotted red line: bursting boundary (50% bursting neurons). Grey in (C, D): parameter combinations where fewer than 15% of neurons burst, with no HFO peak from which to estimate center frequency or PAC. White arrow: the trajectory from the ON operating point (low J, high c̅) to the OFF operating point (high J, low c̅).

Two features of this landscape are central to the clinical interpretation of the model. First, beta oscillations and beta-HFO PAC emerge through a single synchronization transition: the beta-power and PAC boundaries both overlap with the CV = 0.5 boundary (dashed white line in Fig 6B, C). Below the boundary the LFP is spectrally diffuse; above it, relative beta power and PAC rise sharply, with the modulation index significant against circular-shift surrogates (Fig S8, p < 0.05). Second, the HFO center frequency varies continuously across the landscape (Fig 6D) and is set primarily by intrinsic excitability, reflecting the single-neuron relationship between reset potential and intraburst firing rate established in Fig 1C. More negative c̅ produces HFOs in the 200-300 Hz range, and less negative c̅ shifts the center toward 300-400 Hz.

The clinical categorization of STN HFOs into slow (sHFO, 200-300 Hz) and fast (fHFO, 300-400 Hz) components [5,6] thus corresponds to distinct regions of this continuous spectral surface rather than to two distinct oscillatory mechanisms. This interpretation predicts that HFO center frequencies across patients should form a continuum, and that the sHFO/fHFO power ratio reflects the position of the individual patient within the parameter landscape.

### Spectral measures across the synchronization transition

Fig 7 traces the slow oscillation peak frequency as a one-dimensional slice through the (c̅, J) landscape, with coupling strength on the horizontal axis and the curves corresponding to different c̅ values overlaid. The slow oscillation frequency generally decreases with increasing coupling, with the rate of decline and starting frequency depending on c̅. At more negative c̅ values (lower excitability), the slow peak prior to synchronization reflects tonic firing rates and lies above the beta range, dropping sharply into it at the transition. At less negative c̅ values, the slow peak starts within the beta range at weak coupling. For most c̅ values, the slow frequency continues to decline below the beta range at strong coupling, reflecting the lengthening of the interburst interval. This is in contrast to the HFO center frequency (Fig 6D) which is largely determined by c̅, with coupling exerting a secondary modulatory effect, so that HFO peak frequency is primarily stratified by intrinsic excitability. At high excitability (c̅ = -52 mV) the peak sits within the fHFO range (300-400 Hz) across coupling values, while at intermediate excitability (c̅ = -58 mV) the peak enters the sHFO range (200-300 Hz) after the synchronization transition. At low excitability (c̅ = -65 mV) the intraburst frequency remains largely below the conventional HFO range, suggesting that clinically detected HFOs correspond to the intermediate-to-high excitability portion of the parameter landscape. The PAC modulation index (Fig S5) confirms the threshold-like relationship to synchronization: the modulation index (MI) is near zero below the transition and rises steeply above it, with the onset coupling for detectable PAC closely tracking the CV = 0.5 synchronization threshold. Notably, at high excitability the MI curve begins to rise at a lower coupling value and, over that initial range only, rises more gradually, reflecting the fact that partial synchronization of the already-bursting population can produce weak PAC before the full transition is complete.

**Fig 7.**
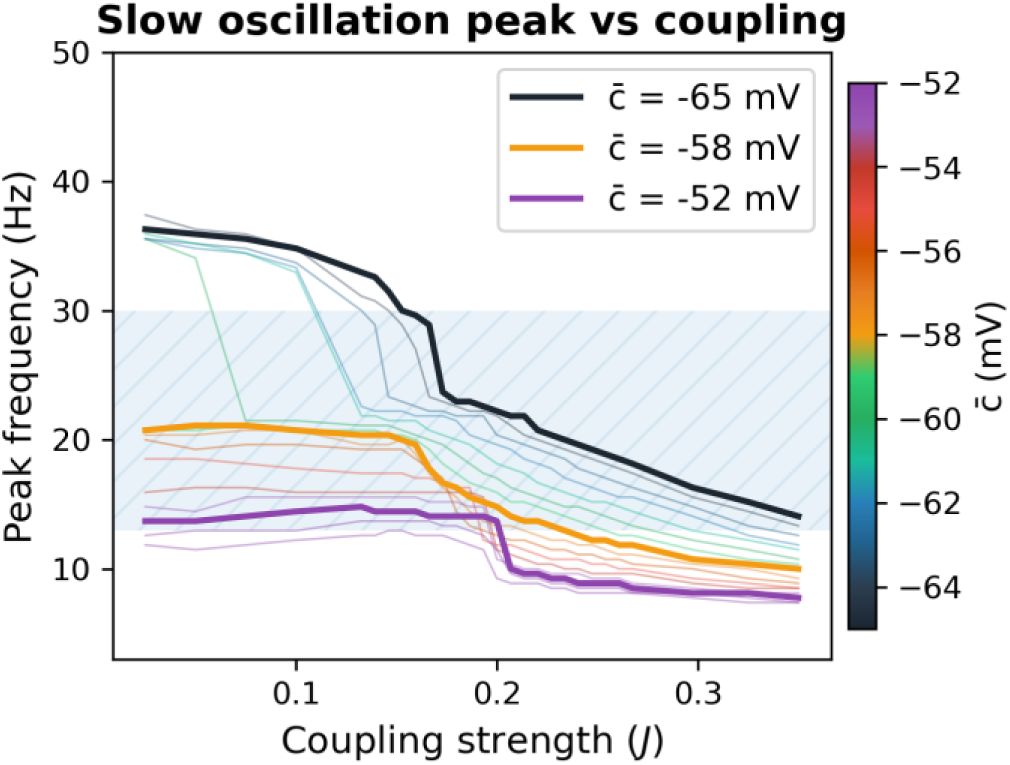
Slow oscillation peak frequency as a function of coupling strength. Curves for different *c̅* values are shown as thin colored lines (color scale at right), with three showcase values highlighted: *c̅* = -65 mV (black), -58 mV (orange), -52 mV (purple). The blue shaded region marks the beta range (13-30 Hz).

### From fast HFO without PAC to slow HFO with PAC: a two-parameter transition

The spectral signatures described above arise from distinct temporal patterns at the population level. Fig 8 illustrates this for two operating points that represent the transition from the medicated-like to the parkinsonian-like state.

**Fig 8.**
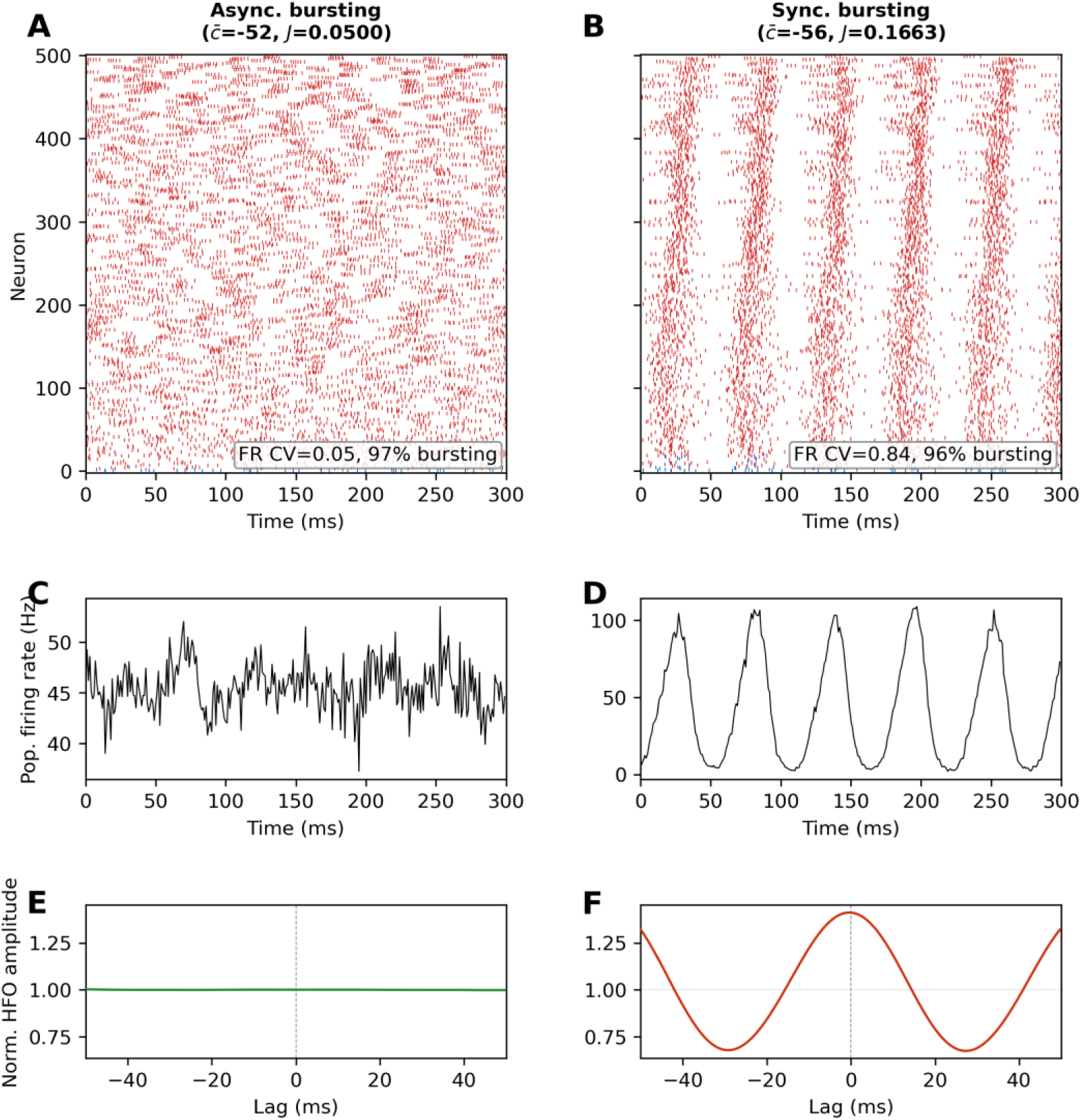
Time-domain dynamics and spike-triggered average HFO envelope amplitude in two regimes. (A, B) Raster plots showing 300 ms of spiking activity in the asynchronous bursting regime (*c̅* = -52 mV, J = 0.050) and the synchronous bursting regime (*c̅* = -56 mV, J = 0.166). (C, D) Mean population firing rate time series. (E, F) Spike-triggered average of the normalized HFO envelope amplitude as a function of lag relative to spike time.

In the first (c̅ = -52 mV, J = 0.050), the population sits in the asynchronous bursting regime: individual neurons burst at high intraburst rates, but burst onsets are uncorrelated across the population (Fig 8A). The resulting mean population firing rate (Fig 8C) fluctuates irregularly, without a clear rhythmic envelope. HFO power in the fHFO range is present but PAC is absent (Fig 9C). In the second (c̅ = -56 mV, J = 0.166), burst onsets are temporally aligned, producing clear population-level volleys visible as vertical stripes in the raster (Fig 8B) and as sharp, periodic peaks in the mean population firing rate (Fig 8D). Here both beta power and beta-HFO PAC are prominent, with the HFO frequency shifting to the lower sHFO range. Mean population firing rates are approximately 44 Hz and 40 Hz in the two regimes, respectively, within the 20-50 Hz range reported for STN neurons in vivo [28].

**Fig 9.**
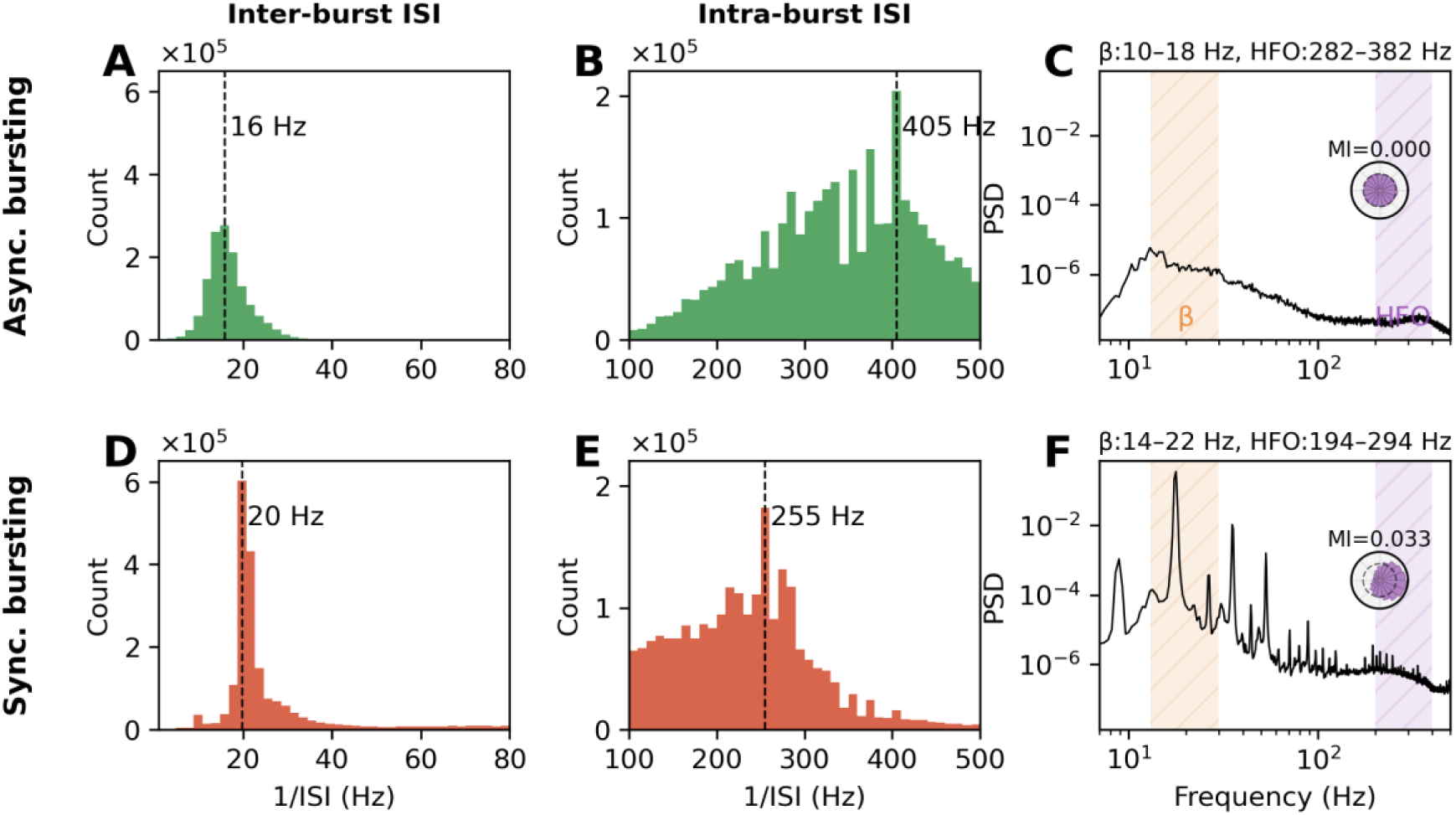
Population ISI frequency distributions and LFP power spectra at two representative operating points. Top row: asynchronous bursting (c̅ = −52 mV, J = 0.050). (A) Inter-burst ISI distribution, peak at 16 Hz. (B) Intra-burst ISI distribution. (C) LFP power spectral density with adaptive PAC bands and modulation index. Bottom row: synchronous bursting (c̅ = −56 mV, J = 0.166). (D) Inter-burst ISI distribution, peak at 20 Hz. (E) Intra-burst ISI distribution. (F) LFP power spectral density with adaptive PAC bands and modulation index. For the non-zero MI value, the surrogate test gives z = 4.0, p < 0.005 (panel F).

Read as the medicated and the parkinsonian states, the transition between these two regimes corresponds to a simultaneous increase in coupling and a decrease in intrinsic excitability. The coupling increase drives the population across the synchronization boundary and produces beta-HFO PAC, while the decrease in excitability lowers the intraburst firing rate and shifts the HFO peak toward the sHFO range, matching the shift toward slower HFOs reported in the parkinsonian relative to the medicated state [5,6]. The decrease in intrinsic excitability may appear to conflict with the higher proportion of burst-firing neurons recorded in the parkinsonian STN [28], since c̅ sets the single-neuron firing mode. The model resolves this tension: the population-level increase in bursting is driven by the coupling increase, not by a change in single-neuron burst propensity. Higher coupling acts as an effective depolarizing drive that recruits neurons across the tonic-to-bursting boundary and, at the same time, aligns their burst timing, so bursting increases even as intrinsic excitability decreases. Because each neuron carries two timescales, fast intraburst spiking and a slow interburst rhythm, this coupling-driven synchronization is a synchronization of bursts, producing the nested beta-HFO oscillation rather than a slow population rhythm alone. The two parameters therefore act on different features of the signal: increasing coupling alone would produce synchronized bursting and PAC but would not shift the HFO frequency, whereas reducing excitability alone would lower the HFO frequency without generating beta-HFO coupling. This provides a possible mechanistic explanation for why the combined HFO ratio and beta power parameter reported by Ozkurt et al. [5] outperforms either biomarker alone: in this framework, each captures a different axis of the underlying parameter space.

The STA of the HFO envelope amplitude (Fig 8E,F) provides a single-neuron view of the same transition. In the asynchronous regime, the STA shows no sustained oscillatory structure, confirming the absence of spike-field coupling. In the synchronous regime, the STA reveals a clear beta-frequency modulation of the LFP locked to individual spike times, consistent with the experimental finding by Meidahl et al. [9] that PAC in the STN reflects spike-field synchronization. The population ISI frequency distributions at these two operating points (Fig 9) confirm that the bimodal temporal structure of individual neuron bursting maps directly onto the two LFP frequency bands: the intraburst ISI peak corresponds to the HFO spectral peak and the interburst ISI peak to the beta spectral peak, with the specific peak frequencies reflecting the different intrinsic excitability at each parameter point, consistent with the single-neuron frequency dependence established in Fig 1C. We note here that the intraburst rhythm is measured directly from the spike trains. Between the two operating points, its peak and the HFO spectral peak both shift down while the beta peak shifts up (Fig 9). This supports the HFO being a separately generated rhythm rather than a beta harmonic, since a harmonic of beta would instead move with the beta peak. The measured coupling is therefore between two distinct rhythms, not an artifact of a non-sinusoidal beta cycle.

### Hysteresis and bistability

We next asked whether this transition could be associated with any hysteresis effects by sweeping the coupling J up and then down while holding the median excitability c̅ at a fixed value (Methods). At c̅ = -62 and -54 mV the up-and down-sweeps follow different branches of steady state synchrony value over a finite interval of coupling strength, so the transition is discontinuous and shows hysteresis, whereas at the intermediate c̅ = -58 mV the two directions coincide and no hysteresis window is detected (Fig 10A-C). The width of the hysteresis window is thus dependent on c̅ and corresponds to a bistability regime in which an asynchronous (Fig 10D) or partially synchronous (Fig 10E) stable state coexists with another strongly synchronized stable state for the same coupling strength (Fig 10D,E).

**Fig 10.**
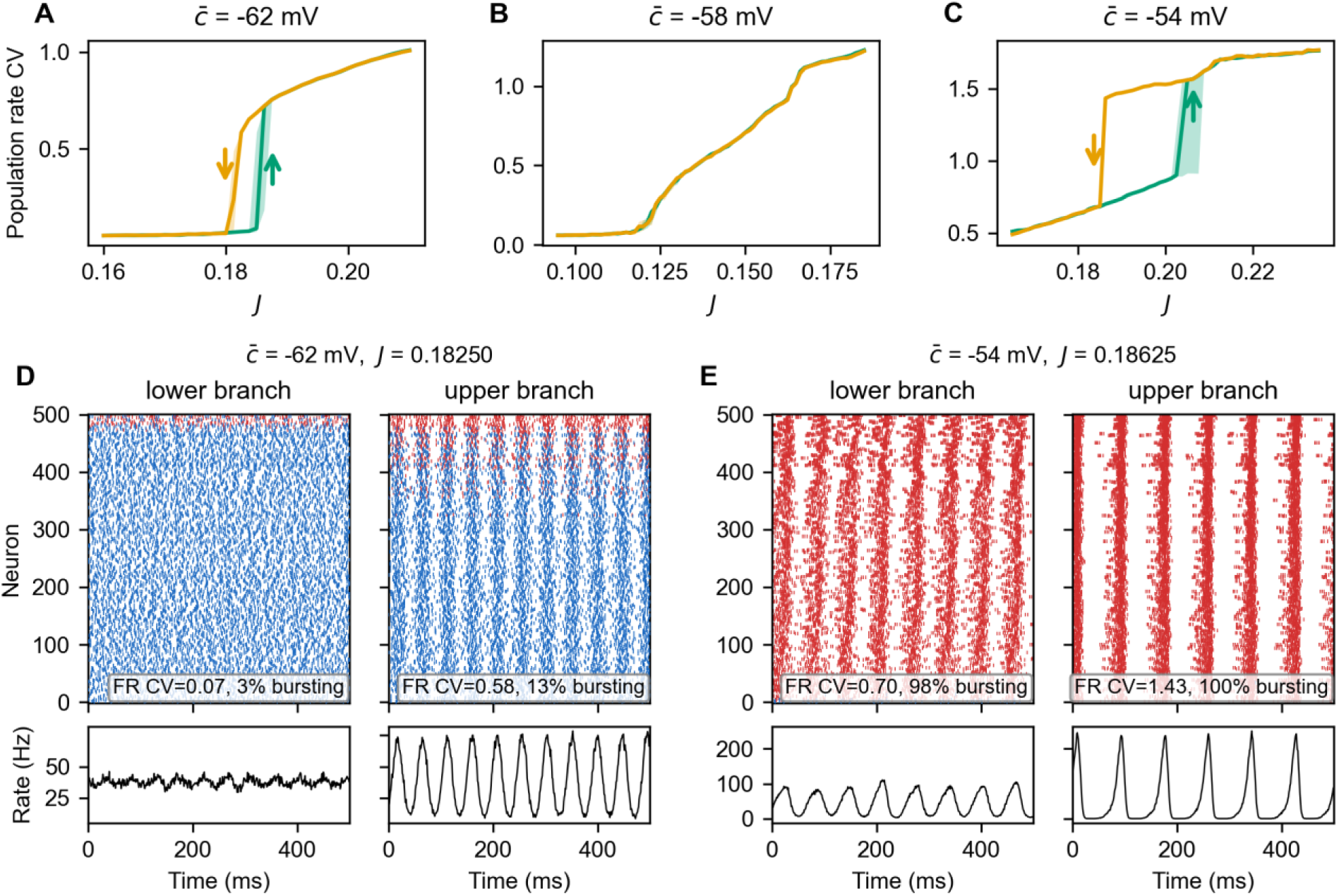
Bistability around the synchronization transition. (A-C) Mean firing rate CV as a function of coupling J for down-sweeps (decreasing J, orange) and up-sweeps (increasing J, bluish green), at three values of median intrinsic excitability: c̅ = -62 mV (A), -58 mV (B), and -54 mV (C). Lines are the median across replications and shaded bands the replication min-max. At c̅ = -62 and -54 mV the up-and down-sweep branches separate over an interval of J, so the synchronization transition shows hysteresis; arrows mark each branch’s jump and its sweep direction. At c̅ = -58 mV the two branches coincide and there is no hysteresis window. (D, E) The two coexisting solutions within the bistability interval, at c̅ = -62 mV, J = 0.18250 (D) and c̅ = -54 mV, J = 0.18625 (E). For each operating point the lower and upper branches are shown side by side: rasters (top; burst spikes red, tonic spikes blue) and the population firing rate (bottom), over the last 500 ms of the steady state. Each raster is annotated with the mean firing rate CV (FR CV) and the percentage of bursting neurons: (D) lower branch asynchronous state, upper branch strong synchrony state; (E) lower branch partial synchrony state, upper branch strong synchrony state.

### The candidate mechanism in an STN-GPe loop

We next examined whether the proposed beta-HFO PAC mechanism could emerge in a circuit that lacks recurrent STN excitation and instead couples the STN to a GPe-like population through a reciprocal excitation-inhibition loop (Methods; Fig S9). The STN reset parameters are recalibrated so that an isolated STN neuron’s interburst rhythm lies well below the beta band (Fig S10), so a beta rhythm in this circuit must arise from the loop rather than from single-cell adaptation. Across the (c̅, J_loop_) landscape the loop reproduces the organization of the single-population model (Fig S11): increasing coupling strength drives the STN population from asynchronous to synchronous bursting, relative beta power and beta-HFO PAC rise together at the synchronization boundary, while the HFO center frequency and intraburst spike frequency remain set by intrinsic excitability. The curves showing the STN synchrony (mean firing rate CV; Fig 11A) and PAC modulation index (Fig 11B) as a function of coupling strength exhibit a similar transition to that seen in the minimal STN model; near zero at low coupling and rising sharply above a coupling strength threshold. Representative operating points trace the STN from asynchronous bursting without PAC to synchronous bursting with beta-HFO PAC (Fig S12), similar to what is observed in the minimal STN model.

**Fig 11.**
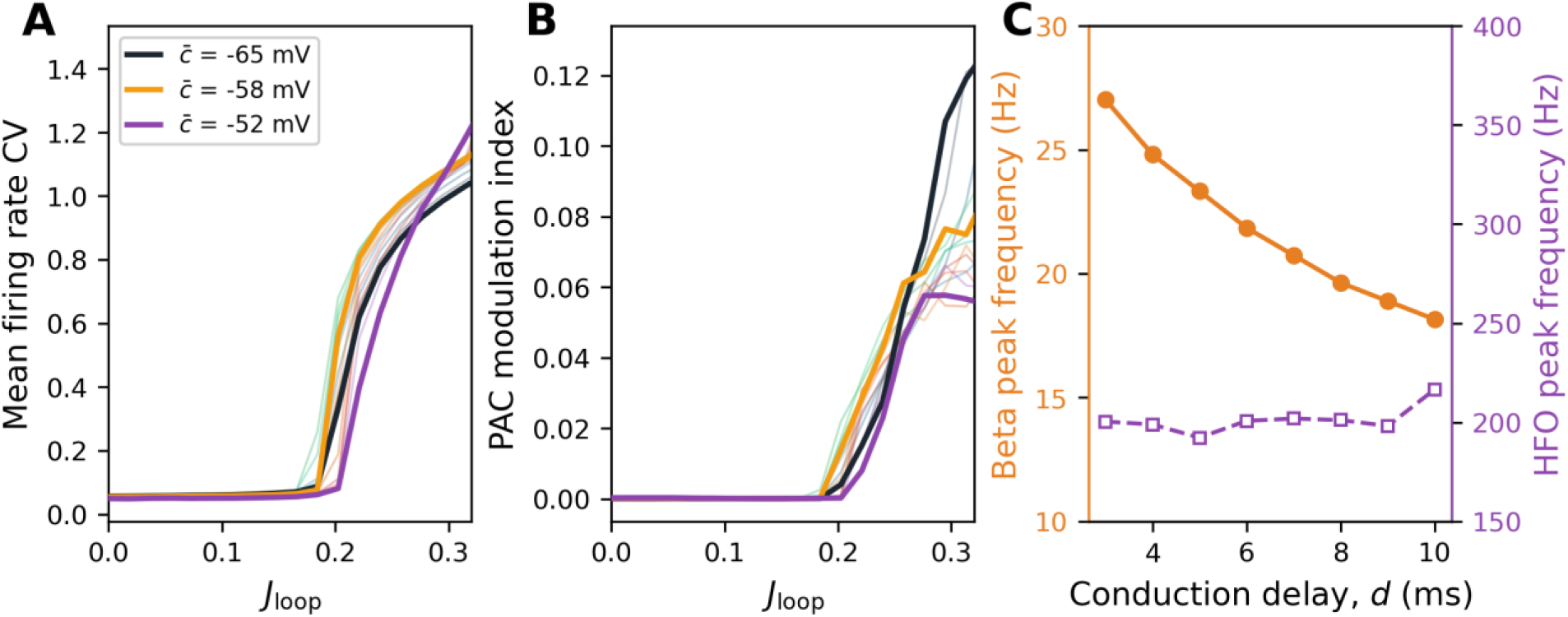
Synchronization, phase-amplitude coupling, and spectral peaks in the STN-GPe loop model. Mean firing rate CV (A) and PAC modulation index (B) of the STN population as a function of the coupling strength J_loop_, at a fixed conduction delay of 6 ms. Each curve corresponds to one value of the median STN excitability c̅; three values are drawn solid and labeled (c̅ = -65, -58, and -52 mV) and the remaining faint color curves follow the same color bar as in Fig 7. (C) Beta peak frequency (left axis) and HFO peak frequency (right axis) of the STN population as a function of conduction delay d, at c̅ = -56 mV and J_loop_ = 0.30.

So the bursting synchronization transition and the emergence of beta-HFO PAC carry over to the STN-GPe loop model, with an important distinction that pertains to the origin of the resulting beta rhythm. The frequency of the beta oscillation is now seen to vary with the value of the delay in the loop synaptic coupling: at a fixed synchronized operating point (c̅ = -56 mV, J_loop_ = 0.30), the beta peak frequency decreases from about 27 Hz to about 18 Hz as the conduction delay increases from 3 to 10 ms, while the HFO frequency remains near 200 Hz and does not track the delay (Fig 11C). Note that the GPe population supplies this synchronizing feedback through rate modulation and does not itself burst (Fig S13), so the HFO and its coupling to beta remain products of the bursting STN population. The proposed mechanism therefore does not require recurrent STN connectivity, and it is consistent with and complementary to classical models in which beta arises from STN-GPe interactions: in this circuit, the loop provides the beta rhythm while the synchronized STN bursting provides the nested HFO and its coupling to the beta phase.

## Discussion

This study presents a minimal spiking network model that provides a unified mechanistic account of the beta-HFO spectral signatures observed in local field potentials from the parkinsonian STN. The central result is that a heterogeneous population of quadratic integrate-and-fire (QIF) spiking neurons with slow adaptation and recurrent excitatory coupling produces three dynamical regimes, tonic firing, asynchronous bursting, and synchronous bursting, governed by two biophysically interpretable parameters. The clinically relevant transition from asynchronous to synchronous bursting reproduces the shift from the medicated to the parkinsonian state. The character of this transition depends on how the population is distributed relative to the single-neuron bursting boundary: it ranges from the co-emergence of bursting and synchrony, through a decoupled transition, to synchronization of pre-existing bursters. The two-parameter landscape yields a continuous spectral surface on which the clinically reported sHFO/fHFO distinction appears as a natural feature of the joint variation of intrinsic excitability (c̅) and synaptic coupling strength (J) rather than evidence for separate oscillatory mechanisms.

### The origin of HFO frequency and medication state dependence

The model provides a potential mechanistic account of what determines HFO frequency. Clinical studies have documented that HFO power shifts from the 200-300 Hz range in the OFF state toward 300-400 Hz in the ON state [4–6], and that the sHFO/fHFO ratio correlates with motor scores [5] and tremor state [29]. However, no previous model has explained what sets the HFO frequency. Our model shows that HFO center peak frequency is the population-level expression of single-neuron intraburst firing rates, which depend continuously on intrinsic excitability and, to a lesser degree, on synaptic coupling strength. This account is consistent with the finding that single-unit power spectra and ISI distributions in human PD STN show peaks at HFO frequencies that correlate with the LFP HFO peak [9], and with the wide dynamic range of STN neurons, which can reach peak firing rates exceeding 200 spikes/s [30]. The sHFO/fHFO distinction reported clinically thus corresponds to sampling different regions of a continuous spectral surface rather than reflecting two discrete oscillatory mechanisms. This interpretation makes a specific prediction: across patients, HFO center peak frequencies should form a continuum rather than clustering into two distinct groups. This follows from the fact that the pathological state, synchronous bursting with beta-HFO PAC, can arise from multiple combinations of intrinsic excitability and coupling strength within the synchronized regime, each producing a different HFO center frequency, a prediction that is testable with large patient cohorts. Coupling of this kind is not exclusive to the STN: enhanced beta-gamma phase-amplitude coupling recorded non-invasively over the scalp also correlates with motor impairment in Parkinson’s disease [31], though the high-frequency component there is broadband gamma rather than the 200-400 Hz STN HFO addressed here.

The two-parameter framework also offers a natural interpretation of the otherwise puzzling finding by Ozkurt et al. [5] that the sHFO/fHFO power ratio correlates with motor scores while beta peak power does not. In our model, HFO frequency varies continuously across the (c̅, J) landscape, while beta power rises steeply at the synchronization transition and saturates in the strongly synchronized regime. These two spectral features are thus partial readouts of different aspects of the underlying parameter space, and their combination better constrains the patient’s overall position in the (c̅, J) landscape, a candidate explanation for why the combined sHFO/fHFO ratio and beta power index reported by Ozkurt et al. [5] outperforms either measure alone. Because the present study doesn’t analyze patient recordings, the Fig 1C relationship of rising HFO frequency with falling beta frequency across excitability is presented as a model prediction rather than an established cross-patient relationship. Testing whether the two peak frequencies covary as predicted across patients remains part of future work.

### The origin of the beta rhythm

In the minimal STN model, the beta-band modulation is set by the interburst interval, which is in turn set by the single-neuron adaptation dynamics: with a recovery timescale of 125 ms and strong post-spike adaptation, the isolated neuron bursts at a beta-range rate (Fig 1C). It should be noted that this employed adaptation timescale is faster than the experimentally reported values for STN neurons [30]; the beta clock in the minimal model should therefore be read as a modeling instrument rather than a biological STN rhythm, and as a different mechanism from the network-level generation of beta through STN-GPe interactions [10,11]. The model, however, does keep beta and HFO as separate rhythms: the interburst interval sets the beta peak while the intraburst firing rate sets the HFO peak (Fig 9), a further indication that the HFO is a distinct rhythm and not a harmonic of beta.

The STN has not been proposed as a sole generator of the beta rhythm; the prevailing view attributes parkinsonian beta to network interactions, in particular the reciprocally coupled STN-GPe circuit together with its cortical and striatal inputs [32]. While the GPe does not itself generate beta, it has been implicated in beta-gamma phase-amplitude coupling when driven by striatal beta input [33]. This motivates the extension of our proposed model to the STN-GPe loop in which beta does not depend on the single-cell adaptation timescale. For that, the STN reset variable is recalibrated so that the isolated interburst rhythm lies well below the beta band (with a corresponding recovery timescale of 500 ms) and there is no recurrent STN excitation, yet the same asynchronous-to-synchronous bursting transition and beta-HFO coupling emerge, with beta generated by the delayed reciprocal feedback between the two populations. This provides an experimentally testable result: a variation in the synaptic delay shifts the beta but not the HFO frequency (Fig 11C), as the latter follows the delay-independent intraburst firing rate. Because excitability and coupling strength affect both beta and HFO frequency, while synaptic delay affects only the beta frequency, this distinction could serve as a model-derived discriminative observation regarding the origin of the beta oscillation.

It is interesting to note that the beta frequency in the examined STN-GPe loop can extend to the higher end of the beta band, whereas for the parameters examined here, the slow peak frequency in the minimal STN model tends towards the lower part of the beta band after the transition (Fig 7). Despite that, both models share the rest of the framework: the synchronization transition, the organization of the spectra by excitability and coupling, and the excitability-set HFO peak frequency.

### Transition scenarios and clinical correlates

The model identifies three qualitatively distinct transition scenarios depending on baseline excitability. HFOs have been reported in the non-parkinsonian human STN [27], suggesting that at least a subpopulation of STN neurons may operate as intrinsic bursters even in the medicated or non-parkinsonian state. However, the proportion of neurons exhibiting burst-like activity is significantly higher in parkinsonian STN compared with non-parkinsonian recordings [28], suggesting that dopamine depletion shifts intrinsic excitability as well as effective coupling. The clinically relevant transition may therefore correspond to the intermediate-excitability scenario, in which dopamine depletion both recruits additional bursters and synchronizes them, with the bursting fraction saturating while synchronization continues to increase (Fig 5B). The medicated state would then correspond to a partially bursting but asynchronous population, where individual neurons produce HFO-frequency intraburst firing but without population-level beta modulation, consistent with the observation that HFOs can be present without pathological PAC in the medicated STN [7,27]. Which scenario applies to a given patient depends on the baseline distribution of intrinsic excitability across the STN population, a quantity that, while characterizable in vitro through slice recordings of STN neurons [21], is not currently measurable in vivo. It is also worth noting that the model predicts that beta-range spectral peaks can emerge without HFO: at low baseline excitability, recurrent coupling slows and partially correlates tonic firing across the population, producing an LFP oscillation in the beta range before neurons transition to bursting (Fig 2B). Together with the observation that HFO can occur without beta modulation, this shows that within the present model, beta and HFO can each arise independently, and beta-HFO PAC emerges specifically when synchronized bursting produces both simultaneously. This is consistent with the view that PAC captures a different or more specific aspect of the pathological state than beta power alone [8]. These spectral features correspond to established subthalamic biomarkers of the parkinsonian state and its treatment, including beta oscillatory activity that tracks clinical improvement [2] and beta-burst dynamics that differ between OFF and ON medication [34], alongside the phase-amplitude coupling marker already noted.

That said, progressive disease likely corresponds to simultaneous movement along both axes of the parameter landscape, since dopamine depletion affects both intrinsic membrane currents [21,22] and synaptic conductances [35,36]. One consequence is that movement along both axes in the parkinsonian direction predicts a downward shift in beta peak frequency (Fig 7), consistent with the finding that beta frequency decreases with dopamine depletion in PD patients [37]. In fact, the model framework predicts that the relative timing and ordering of spectral changes during disease progression, if measurable, could help disentangle the relative contributions of intrinsic and synaptic dopaminergic effects in the STN.

The model’s principal spectral and rate quantities can also be compared directly with recordings from the parkinsonian STN. The beta rhythm peaks at 16-20 Hz across the asynchronous and synchronized states (Fig 9), within the 13-30 Hz beta band of the parkinsonian STN [2,7]. The HFO peak lies in the 200-400 Hz range (Fig 6), overlapping the 150-350 Hz range reported for subthalamic HFO [4,7,9]. The mean firing rate is 40-44 Hz across the two states (Fig 8), within the range recorded for parkinsonian STN units [28]. These values are not independent predictions. The frequencies and the firing rate follow from the model’s timescales and drive, which place single neurons in the bursting regime with interburst intervals in the beta range and intraburst firing in the HFO range. The phase-amplitude coupling is different. It is not tuned, and it emerges only when the population synchronizes, reproducing the trend of beta-HFO PAC being observed in the off-medication STN and its resolution with medication [7,8,9].

The model also connects to a dynamical-systems literature that frames parkinsonian basal-ganglia activity in relation to dynamical transition boundaries and changes in operating points. Prior work places the parkinsonian state near the boundary between synchronized and nonsynchronized dynamics, approached by increased coupling under dopamine depletion [38,39]. Related work maps how firing rate and bursting set the operating regime of the STN-GPe network [40]. These studies address synchronization in the beta band. The present work extends this framing to a second, much faster timescale of observed HFO. Because synchronization here is synchronization of bursts, the same transition that produces the beta rhythm also organizes an HFO an order of magnitude faster, carried by intraburst spiking and coupled to the beta phase. The novelty of the work here relative to that modeling literature is therefore not the proposal of the relevance of a synchronization transition in itself, but in elucidating how a faster HFO oscillation and its coupling to beta could emerge from it, and how the single-neuron firing and collective neuronal modes map onto the spectral features of the corresponding LFP.

A two-directional continuation sweep of the model simulations, as coupling is slowly varied at fixed excitability, has shown that the synchronization transition can exhibit hysteresis, and while the study of the full bifurcation structure around that transition warrants further examination in future work, the reported bistability around the synchronization transition (Fig 10) can offer interesting clinical insight with bearing on response to stimulation interventions. Near the hysteresis window, a desynchronizing input could move the network into the basin of the asynchronous state and produce an effect that outlasts stimulation, reminiscent of carry-over effects reported to occur in the context of deep brain stimulation [41].

Efforts to tailor STN deep brain stimulation to the individual patient increasingly try to leverage the intrinsic electrophysiology of the nucleus. Adaptive deep brain stimulation delivers high-frequency stimulation during periods of elevated beta activity, identified as beta bursts, rather than continuously, and the neuronal dynamics that accompany these bursts have been the object of previous studies [42]. The present model offers an interpretation of what a beta burst could be representing at the level of the single-neuron activity. In the model, elevated beta amplitude reflects population synchronization, during which synchronized bursting generates beta and HFO together. Because the HFO is the intraburst firing and the beta period is the interburst interval, the fast firing during such an episode is confined to a limited portion of each beta cycle rather than spread uniformly across it. In this account a beta burst reflects a nested, two-timescale pattern of synchronized bursting rather than a mere population level increase in firing rate, and resolving this pattern could help specify when and how adaptive stimulation is to be best delivered. The mechanism by which STN stimulation relieves symptoms is itself debated, including whether it acts by reducing subthalamic output [43] or by disrupting abnormal patterns of subthalamic activity [44]; this question is beyond the scope addressed here and while the proposed model contains no stimulation term, the two-timescale structure of the neural patterns it describes suggests that the effect of stimulation could depend on its timing relative to the synchronized bursts, both whether it coincides with a synchronized episode and where within the beta cycle it falls, a prediction that future work could potentially test experimentally.

In summary, the model makes several predictions that clinical and experimental studies could test, and for each we note whether it is specific to the present model or shared with the broader class of synchronization-based accounts. First, HFO center peak frequency should vary continuously across patients rather than separating into slow and fast groups, because the synchronized pathological state can be reached from many combinations of intrinsic excitability and coupling, each with a different intraburst firing rate and therefore a different HFO peak (Fig 6D); this continuum, and its origin in the single-neuron intraburst rate, follows from the present intrinsic-bursting account. Second, beta-HFO phase-amplitude coupling should appear only when bursting is synchronized across the population, so that HFO power without beta-locked amplitude modulation marks an asynchronous bursting regime rather than the absence of bursting (Fig 8, Fig 9); that phase-amplitude coupling requires synchronization is shared with other synchronization-based accounts, whereas the distinct asynchronous bursting regime, with HFO but no PAC, is specific to the present model. Third, moving in the parkinsonian direction along the trajectory marked on the (c̅, J) landscape (Fig 6), a joint increase in coupling and decrease in excitability, predicts a downward shift in beta peak frequency (Fig 7) and an ordered sequence of spectral changes whose relative timing, if measured, could separate intrinsic from synaptic dopaminergic effects; the beta-frequency shift matches existing data, while the ordering prediction is specific to the two-parameter framework. Fourth, if the beta rhythm is generated by network feedback rather than by single-cell dynamics, its frequency should shift with the conduction delay without any shift in the HFO frequency, because the delay sets the slow rhythm and the HFO peak follows the delay-independent intraburst firing rate (Fig 11C). Fifth, if the parkinsonian STN operates near the bistable window identified here, brief desynchronizing stimulation that moves the network into the basin of the asynchronous state could produce an effect that outlasts stimulation, and because the window depends on baseline excitability, any such carry-over should depend on excitability (Fig 10); a lasting effect from bistability is generic to bistable systems, while its dependence on baseline excitability is specific to the present model. Sixth, the beta synchrony targeted by closed-loop and phase-specific stimulation is, in this model, synchrony of bursts rather than of single spikes: each neuron fires an HFO-rate burst per beta cycle, so a pulse locked to a given beta phase is targeting neurons each exhibiting fast HFO spikes within a burst rather than neurons firing a single spike each within the population-aggregated burst. The same phase-locked stimulation could therefore act differently on a burst-synchronized population than on one synchronized at the level of single spikes, a distinction marked by the presence of a genuine nested HFO and its phase coupling and is specific to the present model.

### Relationship to prior computational work

Sanders [19] established the foundational insight that synchronized bursting produces PAC resembling that observed in parkinsonian recordings. Our model extends this in three directions. First, we derive burst statistics as emergent outputs of multiple-timescale neuronal dynamics rather than prescribing them as inputs, enabling the identification of the transition between regimes. Second, we provide a mechanistic account of HFO frequency that was not addressable in their framework. Third, we identify the asynchronous bursting regime (HFO without PAC), which their model treats as the trivial case of zero prescribed synchronization. The Meidahl et al. [9] experimental finding that PAC reflects spike-field synchronization provides direct empirical support for the synchronization-based mechanism common to both works.

Circuit-level models of beta generation [10–14] operate at a different scale and address a complementary question: how oscillations arise from multi-region interactions. Several of these have employed Izhikevich-type neurons in multi-region basal ganglia networks to investigate action selection, deep brain stimulation effects and pathological synchronization [45–47], demonstrating that this neuron class can capture basal ganglia dynamics across scales ranging from small networks to populations of tens of thousands of neurons. These models focus on circuit-level phenomena such as thalamic relay fidelity, beta-band synchrony driven by STN-GPe interactions, and the combined effects of altered intrinsic properties and synaptic conductances following dopamine depletion, but do not address HFO generation or beta-HFO coupling observed in STN local field potentials. Our model uses the same neuron class but exploits the dependence of intrinsic bursting dynamics on the parameters to explain the origin of HFOs and their coupling to beta oscillations, a question that requires resolving the single-neuron firing mode within the population. Our model does not require external beta input; the beta rhythm emerges from synchronized burst timing. However, these accounts are not mutually exclusive. The STN-GPe feedback system has been shown to function as a central pacemaker capable of generating synchronized oscillatory bursting at multiple frequencies [48], and in a full circuit model, cortex-STN-GPe loop dynamics could modulate both intrinsic excitability (through altered cortical drive to STN) and synaptic coupling (through GPe-mediated changes in effective STN coupling [49]).

Conductance-based single-neuron models have characterized STN bursting at slower, empirically observed timescales. Kubota and Rubin showed that a depolarization-activated inward current combined with an NMDA current produces regular bursts below 1 Hz under hyperpolarizing input [50], and a related model reproduced roughly 1 Hz bursts entrained to cortical slow oscillations and strengthened by dopamine depletion, with inhibition supplied by the globus pallidus [51]. These models resolve the ionic basis of STN bursting at the cortical slow-oscillation rate, whereas the present model abstracts from those currents and treats bursting at the beta timescale, where the interburst interval falls in the beta range. The two descriptions target different oscillatory regimes and could offer complementary insight.

### Connection to mean-field theory

As noted in the Introduction, the Izhikevich neuron population with uniform reset potential admits an exact mean-field reduction [16,52–55] that produces mixed-mode oscillations analogous to our beta-HFO pattern, but c drops out of the macroscopic equations in the Ott-Antonsen reduction approach. The mean-field therefore cannot distinguish the macroscopic behavior of a population of intrinsic bursters (less negative c, e.g. −52 mV) and a population of tonic-firing neurons (more negative c, e.g. −65 mV). This is precisely the limitation that matters for the present work: the asynchronous bursting regime, in which individual neurons burst at HFO rates without population coordination, depends on intrinsic excitability (c) being in the range that produces intrinsic bursting, a single-neuron property that is invisible to the macroscopic description provided by known mean-field models. Capturing this regime within a mean-field framework remains an open theoretical challenge and represents a concrete limitation of the mean-field approach for modeling pathological basal ganglia dynamics where the transition between neuronal firing modes is itself the clinically relevant phenomenon.

### Limitations and future directions

Several limitations should be noted. First, the minimal model represents a single excitatory STN population and does not include inhibitory GPe neurons. The STN-GPe system has been shown to constitute a central pacemaker [48], and beta oscillations in PD likely involve STN-GPe reciprocal interactions [10,11] as well as the cortico-subthalamic drive [12,36]. Our single-population approach is partially justified by the experimental finding that isolated STN neurons produce synchronized burst firing through intrinsic glutamatergic axon collaterals without requiring GPe input [23]. We examined an STN-GPe loop as a mechanism control to test whether the results persist as the simplification is relaxed (Fig 11), though this loop remains a reduced circuit without cortical and striatal drive or their dopaminergic modulation. Second, we use the Izhikevich neuron model, which captures qualitative firing mode transitions but does not model specific ionic channels. A conductance-based implementation would enable more direct mapping of dopamine receptor effects onto model parameters and would likely be required for quantitative patient-specific predictions. Third, synaptic interactions in our model use single-exponential AMPA synapses with sparse random connectivity. Incorporating synaptic delays, distance-dependent connectivity, NMDA receptor dynamics, and inhibitory interneurons could modify the transition dynamics, while the richer repertoire of intrinsic ionic currents present in biological STN neurons [56] could support additional dynamical regimes not captured by the two-variable Izhikevich formulation. Fourth, the model does not incorporate the very slow spike frequency adaptation (time constant ∼20 s) mediated by a slowly accumulating K⁺ current that has been characterized in STN neurons [30], which would modulate sustained firing rates and could shape the effective position in the parameter landscape over longer timescales. Finally, the biophysical parameters motivating this model, including the tonic-to-burst transition, intranuclear axon collaterals, and dopaminergic modulation of intrinsic and synaptic properties, are primarily characterized in rodent preparations [21,23,30,35], whereas the clinical spectral signatures we aim to explain are recorded in human patients. This species gap is common to biophysical modeling of the basal ganglia, but quantitative translation to patient-specific predictions will require validation against human single-unit and LFP data.

Additionally, the model produces highly periodic oscillations in the strongly synchronized regime, whereas beta oscillations and beta-HFO PAC in parkinsonian recordings are characteristically non-stationary, appearing as transient bursts of variable duration and amplitude rather than sustained rhythms [34,57]. Although the model exhibits irregular oscillatory dynamics near the synchronization transition, the deeply synchronized regime generates more regular solutions than are typically observed in vivo. Three factors absent from the current model would be expected to introduce realistic temporal variability. First, time-varying cortical and pallidal inputs to the STN would move the effective operating point through the parameter landscape on behaviorally relevant timescales, causing the population to transiently enter and exit the synchronized regime. Second, the slow spike frequency adaptation current characterized in STN neurons [30], which operates on timescales of tens of seconds, could produce ultraslow modulation of effective excitability, driving intermittent transitions between regimes. Third, short-term synaptic depression, which has been documented at glutamatergic synapses within the STN-GPe network [58], would weaken effective coupling during sustained synchronized bursting as synaptic resources deplete, causing the system to fall out of the synchronized regime until recovery restores the coupling strength. This self-limiting mechanism would naturally produce intermittent beta bursts without requiring external modulation, and predicts that burst duration statistics should reflect the interaction between synchronization dynamics and synaptic resource depletion. Incorporating these elements could enable the model to capture the transient beta burst statistics that are increasingly recognized as more informative biomarkers than time-averaged spectral power [34,57]. Beyond these deterministic factors, dynamical noise, absent from the present simulations, provides a further, potentially relevant stochastic mechanism. Fluctuations from channel and synaptic sources, or from variability in the background input, would introduce temporal variability into the oscillations and could shape the resulting beta burst statistics, including burst duration and amplitude. In the bistable regime, the coexistence of asynchronous and synchronized states at the same coupling (Fig 10) offers a specific substrate for this. Noise could then drive intermittent switching between the two states, producing transient rather than sustained coupled beta and HFO. Stochastic simulations lie outside the scope of the present study, so we mention this as an interesting candidate mechanism to be examined in future work rather than a demonstrated one.

Despite these limitations, the model provides an interpretable framework that captures the essential features of STN beta-HFO dynamics and generates specific testable predictions. By dissecting how multiple-timescale mesoscopic LFP spectral features map onto the microscopic neuronal operating regime, the work provides a foundation for future extended models that incorporate circuit-level interactions. More broadly, the ability to infer a patient’s position in the dynamical parameter landscape from their LFP spectral features could inform more effective DBS strategies that target the specific dynamical regime rather than relying on generic spectral biomarker thresholds. For example, patients in the synchronous bursting regime might benefit from desynchronizing protocols such as coordinated reset or phase-specific closed-loop DBS. Because the pathological feature in the model is the synchrony of bursts and not the bursting itself, the aim of such stimulation is to remove the synchrony and restore the asynchronous bursting state of the medicated condition, in which HFO persists without beta modulation, rather than to reduce STN firing rate, per se. Testing these predictions will require extended network models that incorporate GPe inhibition, cortical drive, and their dopaminergic modulation, validated against multi-nucleus recordings in animal models of progressive dopamine depletion. The present work represents a first step in that direction.

## Methods

The neurons are represented by the Izhikevich adaptive quadratic integrate-and-fire model, a reduced phenomenological description rather than a conductance-based STN model. Its parameters are calibrated at the single-neuron level to place the cell in the tonic-to-burster transition and to reproduce the two characteristic timescales of the bursting neuron (Fig 1); the individual parameter values are not intended as direct biophysical measurements. The single-population and STN-GPe loop models share this substrate, and the loop STN reset parameters are recalibrated so that the isolated interburst rhythm lies below the beta band (see STN-GPe loop model).

### Neuron model

We simulate a population of N = 5000 Izhikevich neurons [20]. The membrane potential *v*_*i*_ and recovery variable *u*_*i*_ of each neuron i evolve according to:

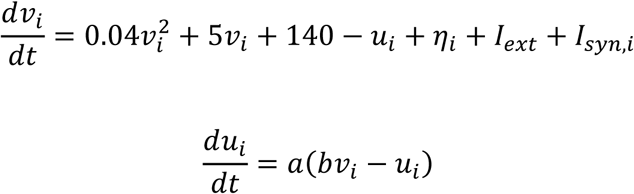

with spike-and-reset rule: when *v*_*i*_ ≥ 30 *mV*, *v*_*i*_ is reset to *c̅*_*i*_and *u*_*i*_is incremented by d. We use parameters a = 0.008, b = 0.2, d = 2.0, and external current *I*_*ext*_ = 8.0. These values place single neurons in a regime where the reset potential *c̅*_*i*_ determines the firing mode: more negative c (∼ -64 mV) produces tonic firing, while less negative c (∼ -52 mV) produces bursting. All model parameters are listed in Table 1.

**Table 1.** Model parameters. Fixed parameters are set to values that place single neurons in the tonic-to-burster transition regime. Swept parameters (*c̅*, J) define the two-dimensional landscape explored in this study.

| Parameter | Symbol | Value | Description |
| --- | --- | --- | --- |
| **NEURON MODEL** |  |  |  |
| Recovery rate | $a$ | 0.008 | Calibrated for beta-range interburst frequency |
| Recovery sensitivity | $b$ | 0.2 | Standard Izhikevich value |
| Recovery increment | $d$ | 2.0 | Controls intraburst spike count and burst termination |
| External current | $I_{ext}$ | 8.0 | Background drive, calibrated for physiological firing rates |
| Spike threshold | $v_{thresh}$ | 30 mV | Standard Izhikevich threshold |
| **POPULATION** |  |  |  |
| Number of neurons | $N$ | 5000 | |
| Reset potential spread | $\Delta_c$ | 1.0 mV | Lorentzian half-width for $c_i$ distribution |
| Background current center | $\bar{\eta}$ | 5.0 | Lorentzian center for $\eta_i$ distribution |
| Background current spread | $\Delta_\eta$ | 2.0 | Lorentzian half-width for $\eta_i$ distribution |
| **CONNECTIVITY** |  |  |  |
| Connection probability | $p$ | 0.2 | Sparse random (Erdős-Rényi) |
| Synaptic time constant | $\tau_{syn}$ | 1.5 ms | Single-exponential AMPA decay |
| Reversal potential | $E_{syn}$ | 0 mV | Excitatory (AMPA) |
| **SIMULATION** |  |  |  |
| **Time step** | $dt$ | 0.05 ms | Half-step Euler integration |
| **Total duration** | $T$ | 23 s | 3 s transient + 20 s analysis |
| **Transient discarded** | $T_{trans}$ | 3 s | |
| **LFP sampling rate** | $f_s$ | 1400 Hz | After downsampling from 20 kHz |
| **Replications per point** | $N_{rep}$ | 12 | Independent random seeds for initial conditions |
| **SWEPT PARAMETERS** |  |  |  |
| **Center reset potential** | $\bar{c}$ | -65 to -50 mV | 16 values, 1 mV steps |
| **Coupling strength** | $J$ | 0.025 to 0.35 | 28 values |
| **ANALYSIS** |  |  |  |
| **Welch window** |  | 3780 samples | ~2.7 s at 1400 Hz, 50% overlap |
| **Mean firing rate CV bins** |  | 2 ms | Spike count histogram bin width |
| **Burst detection: intra-burst** |  | <10 ms ISI | Corresponds to >100 Hz firing |
| **Burst detection: inter-burst** |  | >30 ms ISI | Corresponds to <~33 Hz firing |
| **PAC phase bins** |  | 18 | 20° bins for Tort modulation index |
| **STN-GPe LOOP MODEL** |  |  |  |
| **Number of neurons** | $N_S$ | 5000 | Excitatory (STN) population |
| **Number of neurons** | $N_G$ | 2000 | Inhibitory (GPe) population |
| **Recovery rate** | $a_S$ | 0.002 | STN, recalibrated (500 ms interburst); other STN parameters as in the single-population model |
| **Recovery increment** | $d_S$ | 0.6 | STN post-spike increment |
| **Recovery rate** | $a_G$ | 0.005 | GPe |
| **Recovery sensitivity** | $b_G$ | 0.585 | GPe |
| **Reset potential** | $c_G$ | -65 mV | GPe, homogeneous |
| **Recovery increment** | $d_G$ | 4.0 | GPe post-spike increment |
| **External current** | $I_{ext,G}$ | 28.0 | GPe background drive center |
| **Background current spread** | $\Delta_\xi$ | 2.0 | GPe; Lorentzian half-width, calibrated to 50-70 Hz rate |
| **Reversal potential** | $E_{exc}$ | 0 mV | Excitatory, STN→GPe (AMPA) |
| **Synaptic time constant** | $\tau_{ampa}$ | 2 ms | STN→GPe, AMPA decay |
| **Reversal potential** | $E_{gaba}$ | -80 mV | Inhibitory, GPe→STN and GPe→GPe (GABA-A) |
| **Synaptic time constant** | $\tau_{gaba}$ | 6 ms | GPe→STN and GPe→GPe, GABA-A decay |
| **Loop delay** |  | 6 ms | (STN→GPe and GPe→STN) |
| **Intrapallidal delay** |  | 1 ms | GPe→GPe |
| **Intrapallidal gain** | $W_{gg}$ | 0.10 | Held fixed |
| **Coupling** | $J_{loop}$ | 0 to 0.35 | Swept; 20 values, shared STN-GPe gain |
| **Center reset potential** | $\bar{c}$ | -65 to -52 mV | Loop STN excitability grid; 14 values, 1 mV steps |
| **Loop delay sweep** | | 3 to 10 ms | at $\bar{c} = -56$ mV, $J_{loop} = 0.30$ |

### Population heterogeneity

The reset potential *c̅*_*i*_ for each neuron is drawn from a Lorentzian (Cauchy) distribution centered at c̅ with half-width at half-maximum Δ_*c̅*_ = 1 mV: *c̅*_*i*_∼ Lorentzian(c̅, Δ_*c̅*_). Because the Lorentzian is symmetric, c̅ is the population median: half the neurons have reset potentials above c̅ and half below. The half-width Δ_*c̅*_ = 1 mV corresponds to the interquartile range, so that 50% of neurons fall within c̅ ± 1 mV and the remaining 50% extend further toward the truncated tails of the distribution. This means that the position of c̅ relative to the single-neuron tonic-to-bursting boundary (approximately -57 mV, Fig 1C) directly controls the fraction of the population that bursts intrinsically: when c̅ is well below this boundary, only the upper tail of the distribution produces bursters; when c̅ is above it, the majority of neurons burst. The per-neuron reset potentials are generated from fixed uniform quantiles through the Lorentzian quantile function, with the quantiles restricted to the interval (0.05, 0.95). This truncation bounds the draw to c̅ ± 6.31 mV. The resulting values are then clipped to the fixed interval [-70, -48] mV, which prevents physiologically implausible reset potentials. The external current offsets are drawn through the same quantile truncation and are bounded to [-7.63, 17.63]. In addition, each neuron receives a heterogeneous external current offset drawn from a Lorentzian distribution with center η̅ = 5.0 and half-width *Δ_η_* = 2.0, so that the total external drive to neuron i is *I*_*ext*_ + *η*_*i*_. This additional heterogeneity in external input provides a second source of variability in effective excitability across the population, so that the empirical bursting fraction at a given c̅ reflects the combined effect of both distributions. The Lorentzian form for the reset potentials is a modeling choice, not a requirement of the model, and the population dynamics do not depend on it. Redrawing the reset potentials from a Gaussian of equal full width at half maximum (σ ≈ 0.85 mV) leaves the synchrony, bursting fraction, and phase-amplitude coupling qualitatively similar to those under the Lorentzian (Fig S6). Heterogeneous external input to the STN is itself physiologically motivated by its long-range corticofugal innervation [59], whereas the distributional form of both sources reflects a modeling choice rather than an inference from single-cell recordings. The center reset potential c̅ is varied across 16 values from -65 mV to -50 mV in steps of 1 mV.

### Synaptic coupling

Neurons are coupled through sparse random excitatory connections with connection probability *p* = 0.2. The synaptic current to neuron *i* is conductance-based: *I*_*syn*,*i*_(t) = *g*_*syn*,*i*_(t) · (*E*_*syn*_ − *v*_*i*_(t)), where *E*_*syn*_= 0 mV is the excitatory reversal potential and *g*_*syn*,*i*_(t) is the total synaptic conductance received by neuron *i*, given by: *g*_*syn*,*i*_(t) = (*J*/(*pN*)) ∑ *s*_*j*_(t), where the sum is over presynaptic neurons *j* connected to neuron *i* (determined by the sparse random connectivity with probability *p* = 0.2), *J* is the peak recurrent conductance (the coupling strength), *N* is the population size, and *s*_*j*_(t) is the synaptic gating variable for neuron *j*. The 1/(pN) factor divides the summed input by the expected in-degree, so the mean recurrent drive is set by the peak conductance J and is independent of network size and connection probability. The gating variable obeys *τ*_*syn*_ d*s*_*j*_/dt = −*s*_*j*_, with *s*_*j*_ incremented by 1 at each spike of neuron *j* and *τ*_*syn*_ = 1.5 ms (AMPA kinetics). Note that reducing the connection probability from 0.2 to 0.10 and 0.05 leaves the synchronization transition, and the evolution of the beta-HFO PAC, essentially unchanged (Fig S7). The coupling strength *J* is varied across 28 values from 0.025 to 0.35, sampled with a coarse-fine-coarse density that resolves the synchronization transition. The sparse random topology with *p* = 0.2 balances computational tractability with local connectivity motivated by the compact structure of the STN, where intranuclear axon collaterals have been demonstrated in rodent preparations [23].

### Simulation protocol

The system is integrated using a half-step Euler method (two 0.5*dt substeps per timestep for the voltage equation) with time step dt = 0.05 ms for 23 seconds total simulation time. A Numba-compiled kernel implements the integration loop for computational efficiency. The first 3 seconds are discarded as transient, and all analyses are performed on the remaining 20 seconds. For each (*c̅*, J) parameter combination, 12 independent replications are simulated. The random network realization is fixed: the connectivity matrix and both heterogeneity vectors are constructed once, from a single seed, and are identical across replications. The two heterogeneity sources are drawn from two independent uniform quantile vectors. Only the initial conditions differ between replications (*v*_*i*_ drawn uniformly from [-65, -50] mV, *u*_*i*_ = *bv*_*i*_). Results are reported as means over replications unless otherwise noted.

### LFP proxy

The synthetic LFP is computed as the mean synaptic current across the population: LFP(t) = (1/*N*) ∑ *g*_*syn*,*i*_(t) · (*E*_*syn*_ − *v*_*i*_(t)). This conductance-based proxy captures the aggregate synaptic currents that dominate extracellular field potentials [19,60]. The LFP is downsampled to 1400 Hz for spectral analysis.

### STN-GPe loop model

We consider a two-population STN-GPe loop with no recurrent STN-STN coupling. An excitatory STN population and an inhibitory GPe population are connected by three delayed, sparse, conductance-based projections: an excitatory STN-to-GPe projection with AMPA-like kinetics (reversal potential 0 mV, decay time constant 2 ms) and inhibitory GPe-to-STN and GPe-to-GPe projections with GABAergic kinetics (reversal potential -80 mV, decay time constant 6 ms). Each neuron follows the Izhikevich dynamics defined above, with parameters specific to each population. Synaptic conductances are normalized by the expected in-degree, as in the single-population model. The two reciprocal projections share the coupling strength J_loop_, while the intrapallidal GPe-to-GPe gain is held fixed. Both loop projections carry an effective transmission delay of 6 ms each, and the intrapallidal projection a delay of 1 ms; these are lumped delays that stand for synaptic integration and polysynaptic routing rather than monosynaptic conduction times.

The STN reset parameters are recalibrated so that an isolated STN neuron’s interburst rhythm lies well below the beta band; the recovery-rate parameter is reduced to 0.002, giving a recovery timescale of 500 ms, and the post-spike recovery increment is reduced to 0.6. STN reset potentials and external drive are drawn from Lorentzian distributions using the same truncated-quantile scheme as the single-population model, while GPe neurons use homogeneous intrinsic parameters with a heterogeneous background current calibrated to a prototypical 50-70 Hz firing rate. All STN-GPe loop parameters are listed in Table 1. Because the loop has no recurrent STN-STN synapse, the LFP proxy is the population summed total membrane current rather than the summed synaptic current; its absolute scale does not affect spectral peak frequencies or relative band powers. The loop is simulated with 5000 STN and 2000 GPe neurons at connection probability 0.2, over the same 23 s protocol and the same fixed random network realization and replication scheme as the single-population model. The excitability-coupling grid spans 14 values of c̅ from -65 to -52 mV, with the coupling strength J_loop_ taking 20 values from 0 to 0.35. In a separate one-dimensional sweep, the loop delay is varied from 3 to 10 ms at c̅ = -56 mV and J_loop_ = 0.30 to test whether the beta peak tracks the delay.

### Spectral analysis

Power spectral density is estimated using Welch’s method with ∼2.7-second Hanning windows (3780 samples at 1400 Hz), 50% overlap, and per-segment linear detrending, yielding ∼0.37 Hz frequency resolution. Relative beta power is computed as the integral of the PSD in the 13-30 Hz band divided by the total spectral power. The HFO center peak frequency is detected on the replication-mean PSD. A minimum filter with a 40 Hz window extracts the local spectral floor, which is smoothed with a 30 Hz Gaussian kernel; the most prominent floor maximum in the 100-600 Hz band, subject to a 50 Hz minimum separation, is taken as the HFO center frequency. Because the minimum-filter window exceeds the 20-25 Hz spacing of the beta harmonics, narrow harmonic lines are removed from the floor and cannot be returned as the HFO peak. A floor maximum is detected only if its prominence is at least 2% of the largest floor value above 150 Hz; when more than one floor local maximum exists, the HFO frequency is taken to be the power-weighted PSD centroid over 200-500 Hz. The same floor detector is applied per replication to identify the adaptive amplitude frequency band used for the modulation index and the spike-triggered average computations.

### Slow oscillation peak detection

To characterize the fundamental frequency of the slow oscillation, the mean PSD across replications is smoothed with a Gaussian kernel (sigma = 2 Hz) in the 3-50 Hz range. Peaks are identified with minimum separation of 3 Hz and minimum prominence of 0.05 (relative to the smoothed PSD range). The lowest-frequency prominent peak is taken as the fundamental slow oscillation frequency.

### Phase-amplitude coupling

PAC is quantified using the Tort modulation index (MI) [61]. The phase frequency band is centered on the detected slow oscillation peak +/-4 Hz. The amplitude frequency band is centered on the HFO spectral peak +/-50 Hz, with a lower bound floor of 80 Hz to prevent overlap with the phase band. If no slow peak is detected (i.e., in the asynchronous regime), a fallback scan over 8-35 Hz phase frequencies is performed to detect any residual coupling. This PSD-guided approach ensures that PAC is computed at the frequencies where oscillatory power actually exists, avoiding the sensitivity to arbitrary band definitions that can affect fixed-band analyses.

PAC is computed on the full 20-second post-transient LFP segment, matching the window used for all other spectral analyses. The phase signal is obtained by bandpass filtering and Hilbert transform, and the amplitude signal is obtained similarly in the HFO band. The phase is divided into 18 bins of 20 degrees each, and MI is computed as the Kullback-Leibler divergence of the amplitude distribution from the uniform distribution, normalized by log(18).

Every reported modulation index is accompanied by a surrogate significance test. For each replication, a null distribution of 200 surrogate modulation indices is generated by circularly shifting the HFO amplitude envelope relative to the beta phase by a random offset drawn uniformly between one tenth and nine tenths of the record length, and recomputing the modulation index on the same bands. The observed modulation index is then reported with the corresponding z-score and a p-value relative to this null distribution. Note that the z-score is partly skewed towards smaller values as the periodicity of the signal increases in the high synchrony regime; this is due to the fact that a time-shift null probes only the alignment between the beta phase and the HFO envelope, so it weakens in the deeply synchronized regime, where the signal is nearly periodic and a shifted envelope can remain aligned with the phase at multiples of the oscillation period.

### Burst detection and classification

Individual neurons are classified as bursting or tonic based on their ISI distributions. A neuron is classified as bursting if it has at least one ISI below 10 ms (intraburst, corresponding to >100 Hz firing) and at least one ISI above 30 ms (interburst, corresponding to <∼33 Hz), indicating a bimodal ISI structure. This dual-threshold criterion provides a simple, parameter-free separation of burst and tonic neurons without requiring histogram fitting. The percentage of bursting neurons at each (*c̅*, J) point is reported as a population-level summary measure.

The intraburst firing frequency of a neuron is also used as a spike-based measure. Interspike intervals in the 1 to 10 ms range are pooled across all neurons and binned at 0.25 ms, and the most common interval is converted to a frequency in hertz as 1000 divided by the interval in milliseconds.

In the STN-GPe loop, where GPe and STN neurons are compared on the same footing, a modified composite classifier is applied to account for the higher firing rate of GPe neurons. A neuron is counted as a burster if three conditions hold: its distribution of log-transformed interspike intervals is bimodal by the Hartigan dip test (p < 0.05); an exact one-dimensional two-means split of those log intervals separates a fast and a slow group whose median-interval ratio is at least 2.3; and at least one burst event is present, defined as a run of intervals below the split threshold, containing at least three spikes. The same thresholds apply to both populations as the ratio criterion is scale-invariant.

### Synchronization measure

Population synchrony is quantified by the coefficient of variation (CV) of the population spike count (computed in 2 ms bins). High CV indicates rhythmic, synchronized population activity (bursts of coordinated firing alternating with quiescence), while low CV indicates asynchronous firing with approximately constant population rate. The synchronization boundary is defined as the (*c̅*, J) contour where CV = 0.5.

### Spike-triggered average HFO analysis

To compare with the experimental findings of Meidahl et al. [9], we compute the STA of the HFO amplitude. The LFP is bandpass filtered in a ±50 Hz window centered on the detected HFO peak frequency (with a lower bound of 80 Hz), and the analytic amplitude (envelope) is extracted via the Hilbert transform. For each spike of every neuron emitting at least 10 spikes in the measurement window, the HFO amplitude in a +/-50 ms window around the spike time is extracted. The mean across all spike-triggered segments is reported. This analysis is performed separately in the asynchronous and synchronous bursting regimes to test whether the model reproduces the experimentally observed distinction between recording states with and without PAC detected.

### Bistability continuation

To test for path dependence in the synchronization transition, the coupling J is swept down and then up at fixed c̅ over a common grid, carrying the full network state continuously from one grid step to the next. The down-sweep carries the synchronized state toward lower coupling and the up-sweep carries the desynchronized state toward higher coupling. Because the stored synaptic conductance scales with J, at each step the conductance is rescaled by the ratio of the new coupling to the old, so that the physical synaptic gating state is preserved; a numerical check confirms that the continuation kernel reproduces the pinned model exactly. The sweeps are deterministic on the single random network realization. At each step the first 3 s are discarded and synchrony is measured over the following 7 s, with an additional 9 s discarded at the cold-started first step of each chain. A survey pass at 0.005 spacing locates the up-and down-jumps in the synchrony measure, a refinement pass at one quarter of that spacing zooms in around the jumps, and a final pass repeats the refinement grid with the per-step transient tripled to confirm that each branch has settled. Hysteresis at a given c̅ is reported when the up-sweep jump bracket lies entirely above the down-sweep jump bracket in every replication. The continuation is run at N = 5000 with four replications and repeated across c̅ to map how the bistable interval depends on excitability.

## Acknowledgments

HS would like to thank Theoden I Netoff for insightful discussions during this work.

## Author Contributions

**Conceptualization**: HS, LAJ, JW, JEA, JLV. **Methodology**: HS. **Software**: HS. **Formal Analysis**: HS. **Investigation**: HS. **Visualization**: HS. **Writing -Original Draft**: HS. **Writing -Review & Editing**: HS, LAJ, JW, JEA, JLV. **Resources**: LAJ, JW, JEA, JLV. **Funding Acquisition**: HS, JLV.

## Funding

This work was supported by the University of Minnesota’s MnDRIVE (Minnesota’s Discovery, Research and Innovation Economy) initiative. This work was also supported by the Udall Center of Excellence for Parkinson’s Disease Research, National Institutes of Health, National Institute of Neurological Disorders and Stroke (P50-NS123109).

## Competing Interests

The authors have declared that no competing interests exist.

## Data and Code Availability

This study is based entirely on computational simulations and does not involve experimental data. All simulation code, analysis scripts, and figure-generation code are publicly available at a Zenodo repository (https://doi.org/10.5281/zenodo.22254023).

## Supporting information

**S1 Fig.**
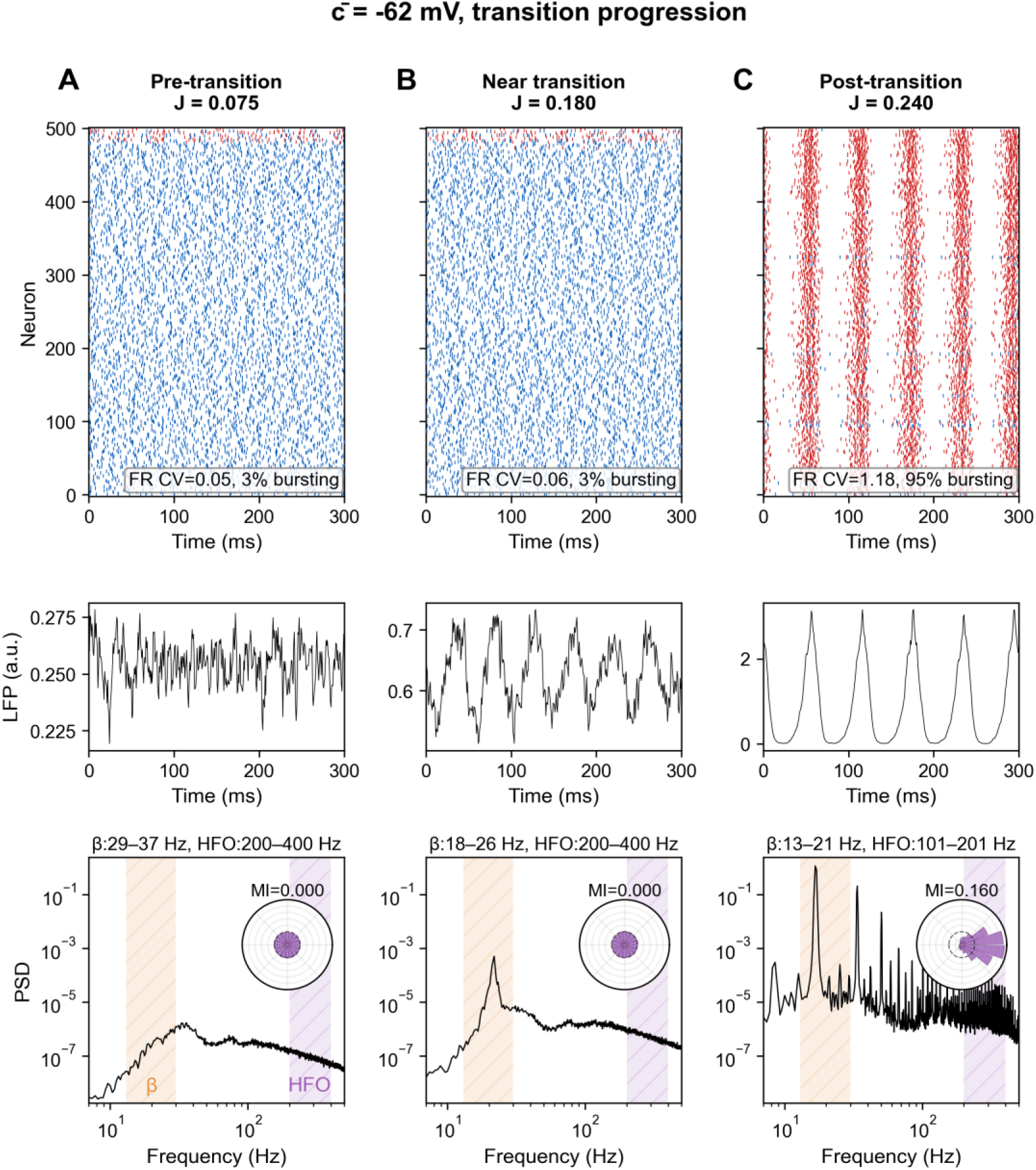
Transition progression at *c̅* = -62 mV. Three operating points: pre-transition (A, J = 0.075), near-transition (B, J = 0.180), and post-transition (C, J = 0.240). For the non-zero MI value, the surrogate test gives z = 2.1, p < 0.05 (panel C). Format as in Figs 2–4.

**S2 Fig.**
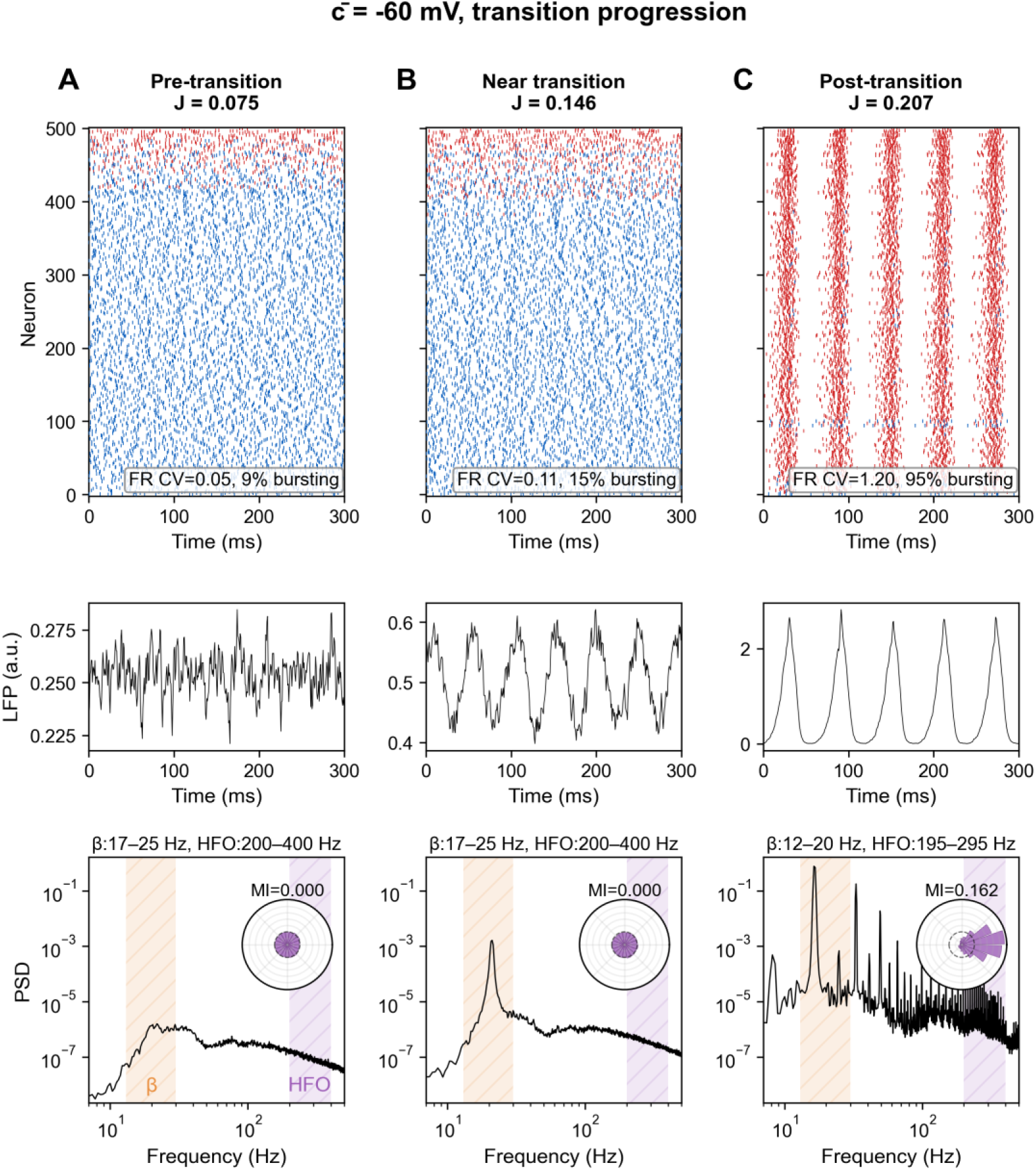
Transition progression at *c̅* = -60 mV. Three operating points: pre-transition (A, J = 0.075), near-transition (B, J = 0.146), and post-transition (C, J = 0.207). For the non-zero MI value, the surrogate test gives z = 4.4, p < 0.005 (panel C). Format as in Figs 2–4.

**S3 Fig.**
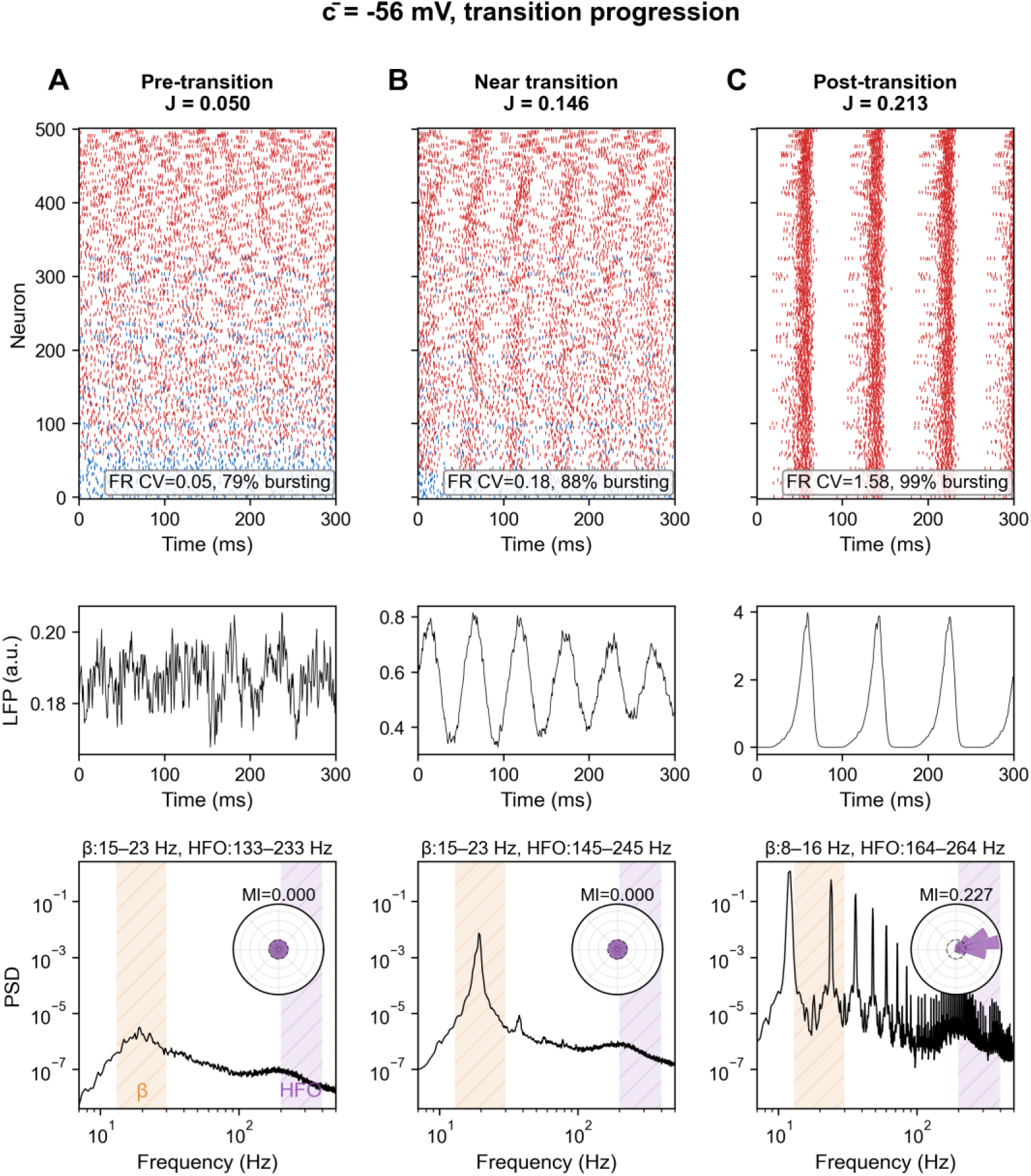
Transition progression at *c̅* = -56 mV. Three operating points: pre-transition (A, J = 0.050), near-transition (B, J = 0.146), and post-transition (C, J = 0.213). For the non-zero MI value, the surrogate test gives z = 1.3, p = 0.11 (panel C). Format as in Figs 2–4.

**S4 Fig.**
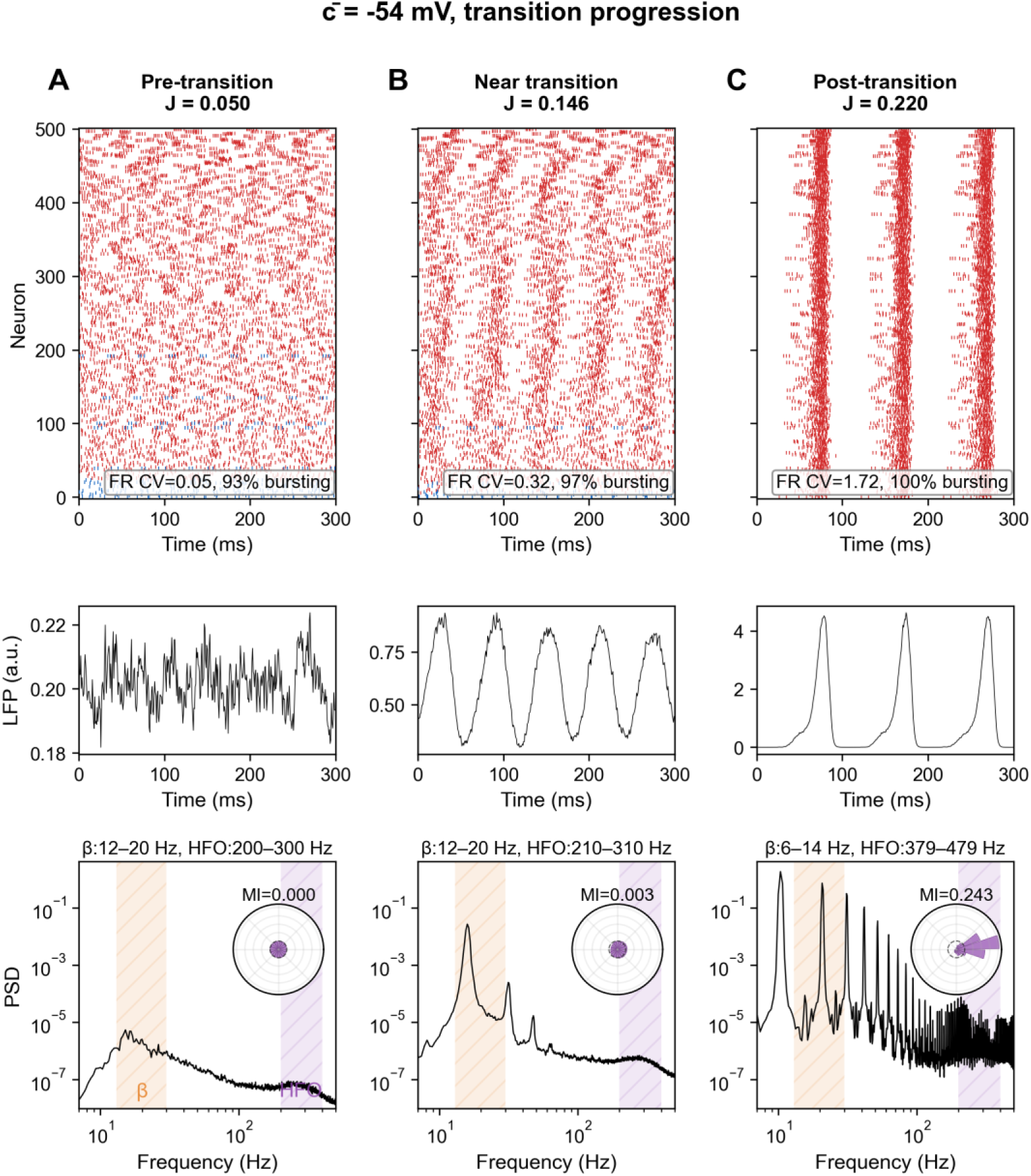
Transition progression at *c̅* = -54 mV. Three operating points: pre-transition (A, J = 0.050), near-transition (B, J = 0.146), and post-transition (C, J = 0.220). For the non-zero MI values, the surrogate test gives z = 17.7, p < 0.005 (panel B) and z = 4.5, p < 0.005 (panel C). Format as in Figs 2–4.

**S5 Fig.**
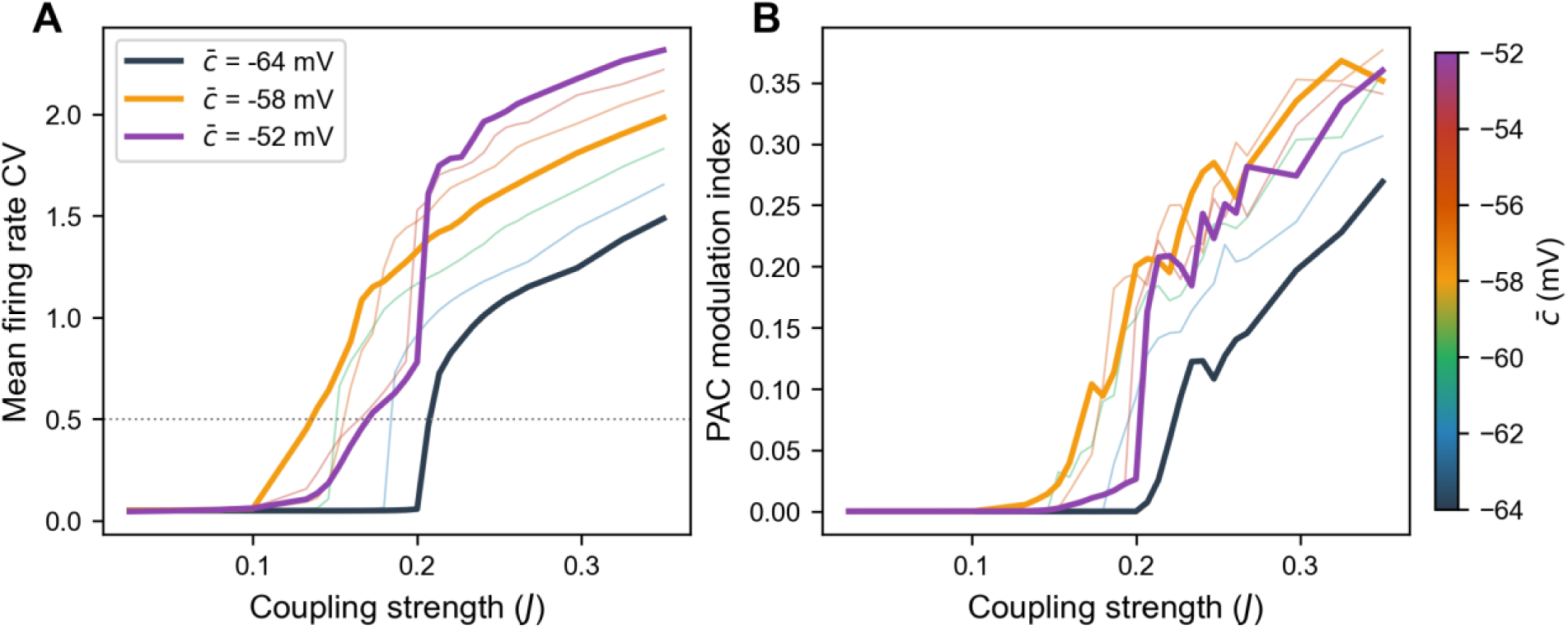
Mean firing rate CV and PAC modulation index versus coupling. (A) Mean firing rate CV and (B) PAC modulation index versus coupling strength J. Thin colored lines show c̅ values from -64 to -52 mV in 2 mV steps (color scale at right); three showcase values are highlighted (c̅ = -64 mV, black; c̅ = -58 mV, orange; c̅ = -52 mV, purple). The dotted line in (A) marks the synchronization threshold (CV = 0.5).

**S6 Fig.**
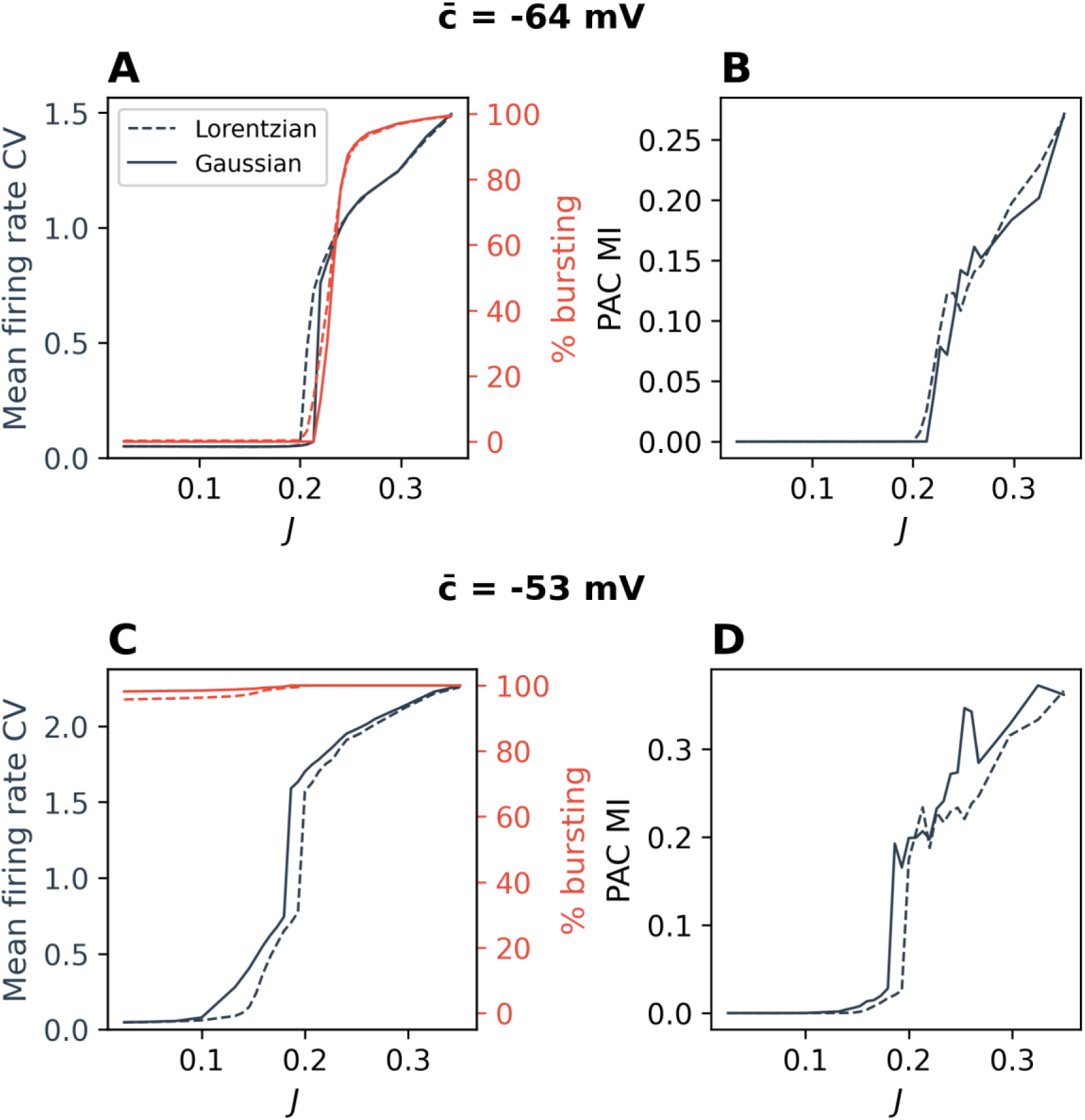
Lorentzian versus Gaussian reset-potential distribution. Reset potentials drawn from the Lorentzian used throughout (dashed) versus a Gaussian of equal full width at half maximum (solid), at two excitabilities (c̅ = -64 mV, top; c̅ = -53 mV, bottom). (A, C) Mean firing rate CV (left axis) and percent bursting neurons (right axis, red) versus coupling J. (B, D) PAC modulation index versus J.

**S7 Fig.**
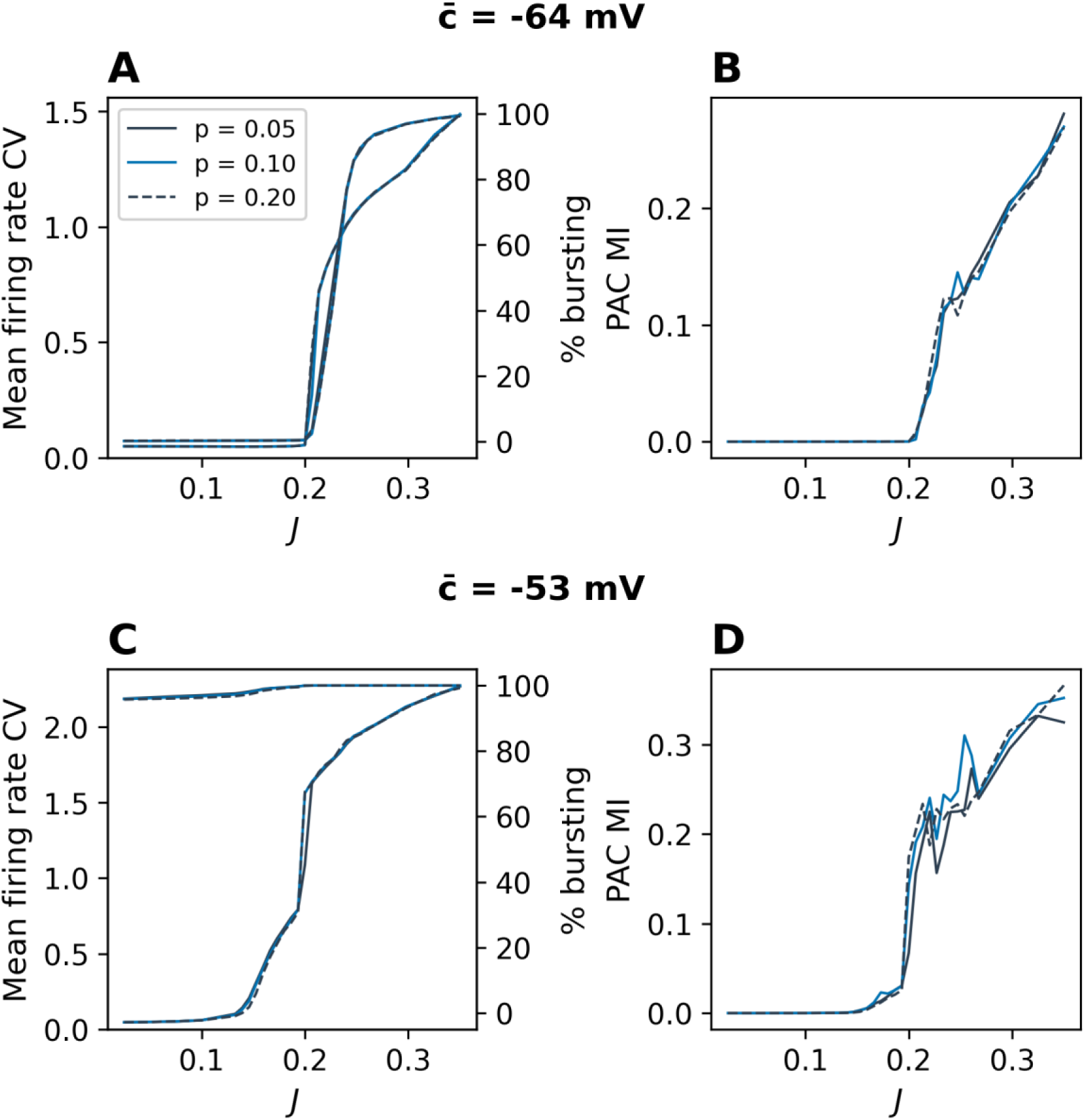
Connection probability comparison. Three connection probabilities (p = 0.05, p = 0.10, and the baseline p = 0.20, dashed), at two excitabilities (c̅ = -64 mV, top; c̅ = -53 mV, bottom). (A, C) Mean firing rate CV (left axis) and percent bursting neurons (right axis) versus coupling J. (B, PAC modulation index versus J.

**S8 Fig.**
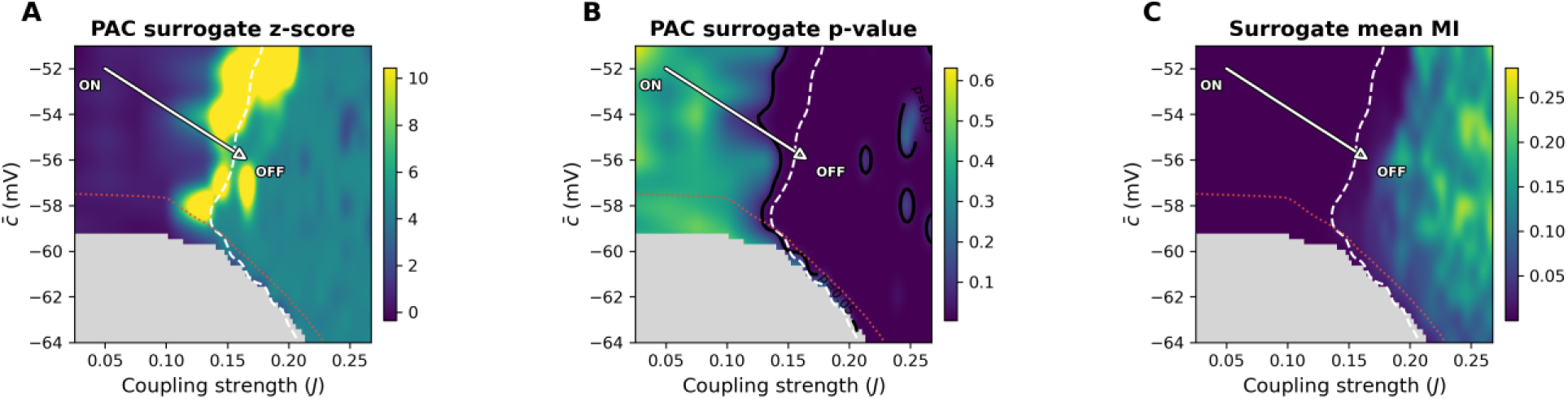
PAC surrogate statistics across the phase diagram. (A) PAC surrogate z-score (color capped at the 90th percentile), (B) surrogate p-value, and (C) surrogate mean modulation index across the (c̅, J) landscape; A and B are replication means. The null is a circular-shift surrogate distribution (200 surrogates). The black contour in (B) marks p = 0.05. Dashed white line: synchronization boundary (CV = 0.5). Dotted red line: bursting boundary (50%). Grey: fewer than 15% of neurons burst or no HFO peak detected. White arrow: the ON-to-OFF trajectory as in Fig 6.

**S9 Fig.**
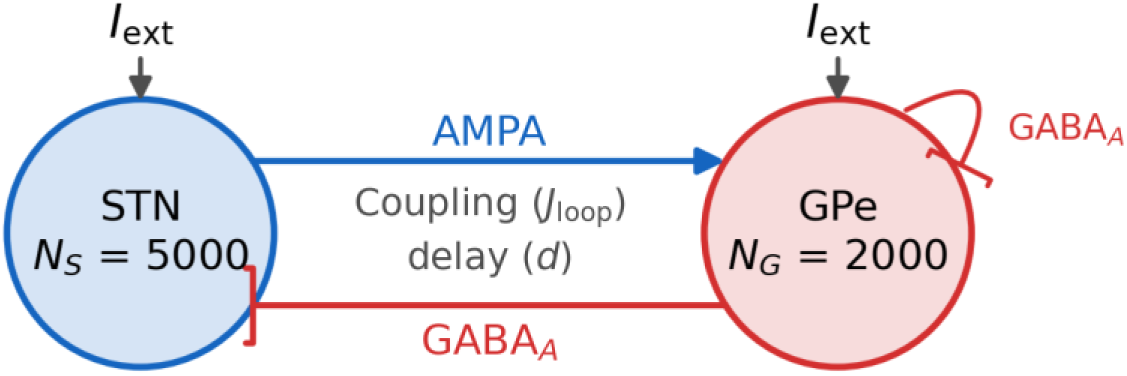
STN-GPe loop circuit. Two-population STN-GPe loop: an excitatory STN population (N_S_ = 5000) and an inhibitory GPe population (N_G_ = 2000), each receiving external drive I_ext_. STN-to-GPe is an excitatory AMPA projection; GPe-to-STN and recurrent GPe-to-GPe are inhibitory GABA_A_ projections. The two reciprocal projections share the coupling strength J_loop_ and a per-leg delay d; there is no recurrent STN-STN coupling.

**S10 Fig.**
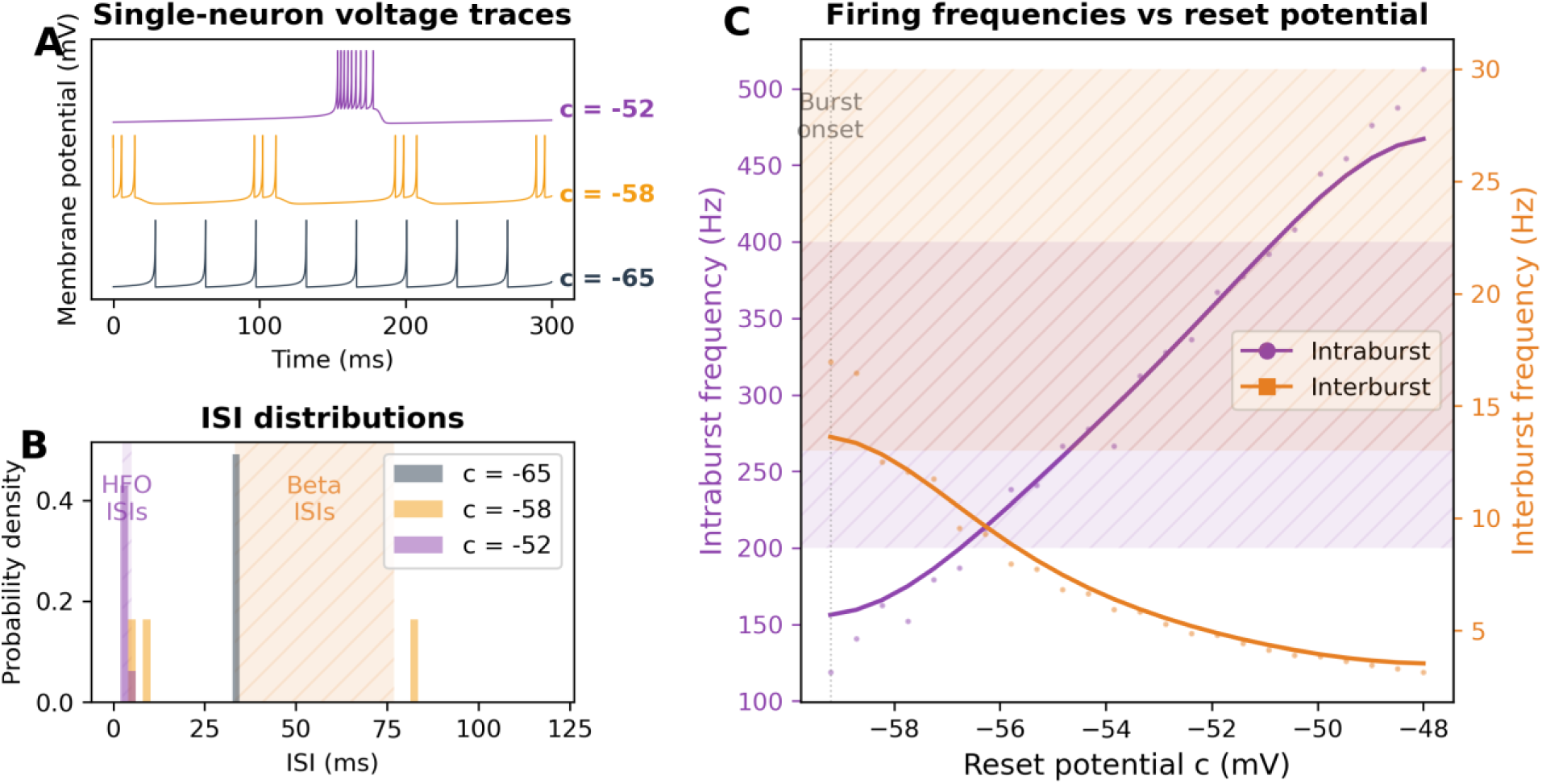
STN-GPe Loop-model STN single-neuron phenomenology. Single-neuron dynamics of the recalibrated STN cell used in the STN-GPe loop model, presented as in Fig 1. (A) Voltage traces at representative reset potentials: tonic (c = -65 mV), short bursts (c = -58 mV), robust bursting (c = -52 mV). (B) ISI probability density distributions at the same reset potentials; shaded bands mark the HFO (purple) and beta (orange) ISI ranges for reference. (C) Intraburst (purple, left axis) and interburst (orange, right axis) frequencies as a function of reset potential c; dots show values at each c, solid lines are smoothed trends, and the dotted vertical line marks burst onset.

**S11 Fig.**
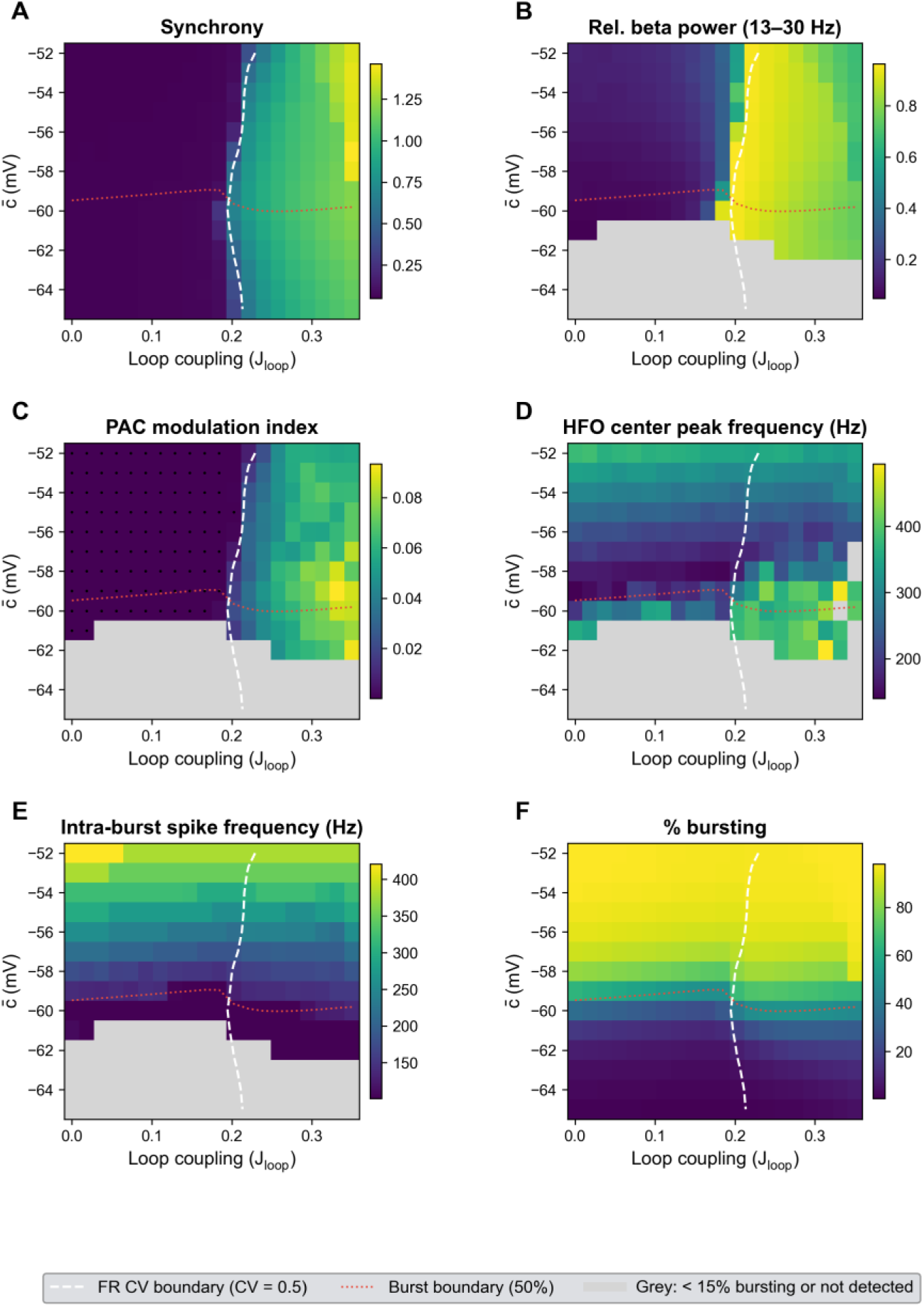
STN-GPe loop phase diagram across the (c̅, J_loop_) landscape. (A) Synchrony (mean firing rate CV). (B) Relative beta power (13-30 Hz). (C) PAC modulation index. (D) HFO center peak frequency. (E) Intra-burst spike frequency. (F) Percent bursting. Dashed white line: synchronization boundary (CV = 0.5). Dotted red line: bursting boundary (50%). Grey: fewer than 15% of neurons burst or no peak detected.

**S12 Fig.**
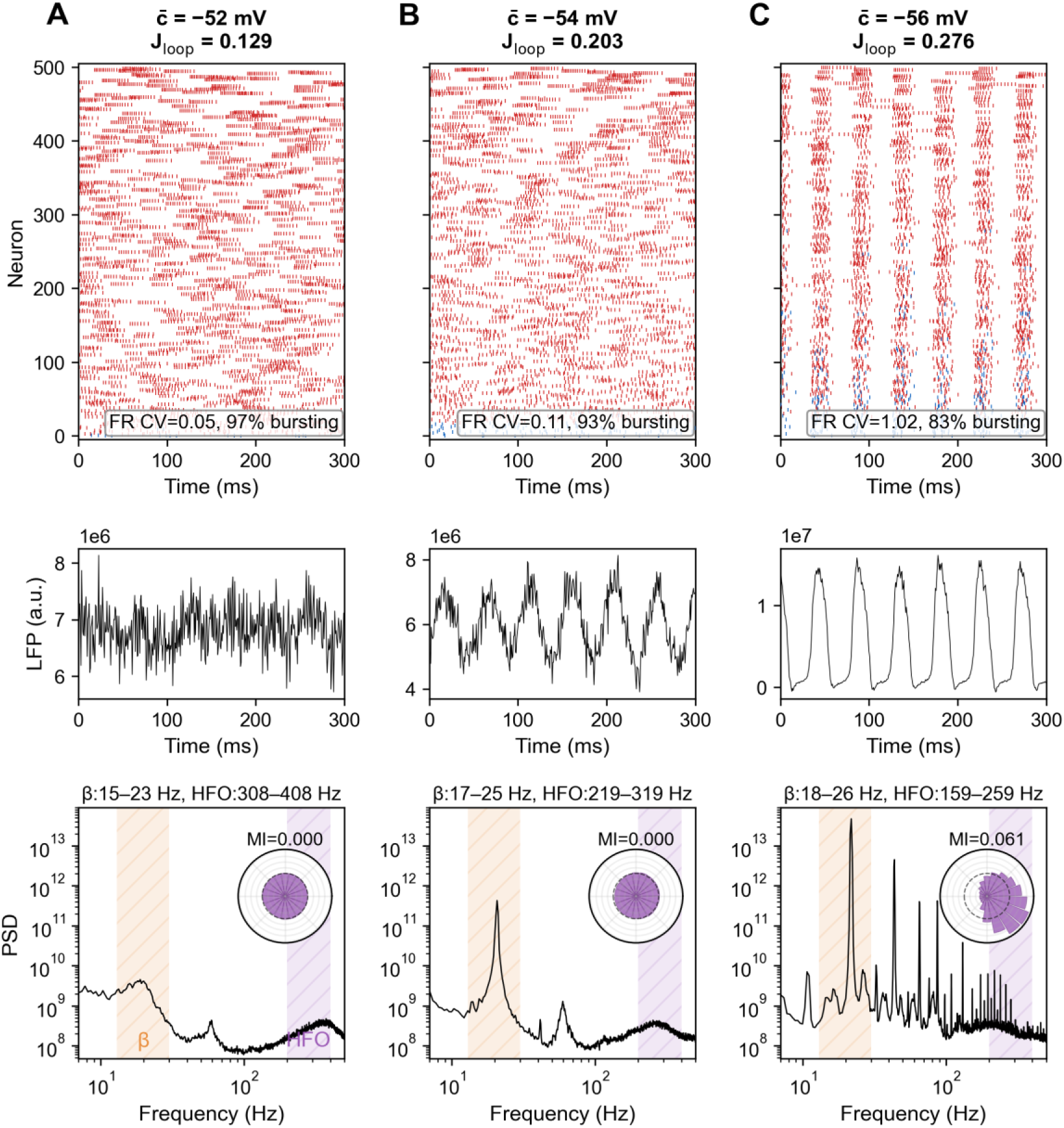
STN-GPe Loop route, STN population. Three operating points: (A) c̅ = -52 mV, J_loop_ = 0.129; (B) c̅ = -54 mV, J_loop_ = 0.203; (C) c̅ = -56 mV, J_loop_ = 0.276. For the non-zero MI value, the surrogate test gives z = 4.3, p < 0.005 (panel C). Format as in Figs 2–4.

**S13 Fig.**
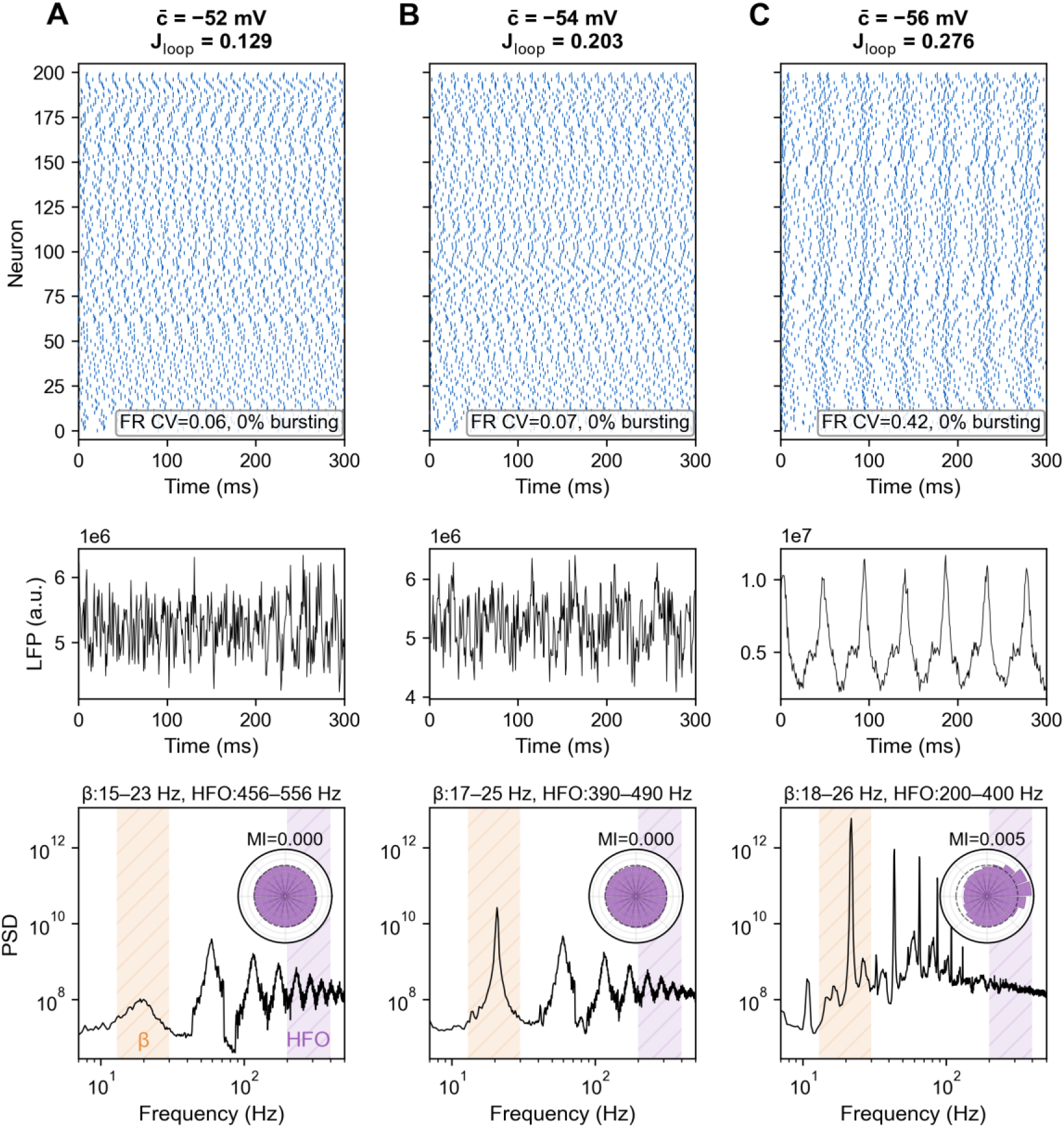
STN-GPe Loop route, GPe population. The GPe companion to Fig S12 at the same three operating points: (A) c̅ = -52 mV, J_loop_ = 0.129; (B) c̅ = -54 mV, J_loop_ = 0.203; (C) c̅ = -56 mV, J_loop_ = 0.276. For the non-zero MI value, the surrogate test gives z = 4.1, p < 0.005 (panel C). Format as in Figs 2–4.

## Notes

### Competing Interest Statement

The authors have declared no competing interest.

### Summary of Updates

This version has been revised in response to peer review. The principal new material is an extension of the single-population model to an STN-GPe loop, in which the STN is coupled to a pallidal inhibitory population through a delayed reciprocal excitation-inhibition loop with no recurrent STN excitation, and which reproduces the same asynchronous-to-synchronous bursting transition and beta-HFO coupling. A new analysis reports bistability and hysteresis at the synchronization transition in the minimal STN model. Robustness checks for reduced recurrent connectivity and for the choice of heterogeneity distribution have been added, together with surrogate-based significance testing of the phase-amplitude coupling. The Discussion has been expanded with sections on the origin of the beta rhythm, the relation to prior conductance-based bursting models, clinical correlates and deep brain stimulation, and the model's testable predictions. New supplementary figures accompany these additions, and all simulation, analysis, and figure code has been deposited publicly.

